# Mediator 1 ablation induces enamel-to-hair lineage conversion through enhancer dynamics

**DOI:** 10.1101/2022.09.08.507153

**Authors:** Roman Thaler, Keigo Yoshizaki, Thai Nguyen, Satoshi Fukumoto, Pamela Den Besten, Daniel D. Bikle, Yuko Oda

**Affiliations:** Department of Orthopedic Surgery, Mayo Clinic, Rochester, MN, USA; Center for Regenerative Medicine, Mayo Clinic, Rochester, MN, USA; Section of Orthodontics and Dentofacial Orthopedics, Division of Oral Health, Growth and Development, Kyushu University Faculty of Dental Science, Fukuoka, Japan; Section of Pediatric Dentistry, Division of Oral Health, Growth and Development, Kyushu University Faculty of Dental Science, Fukuoka, Japan; Division of Pediatric Dentistry, Department of Oral Health and Development Sciences, Tohoku University Graduate School of Dentistry, Sendai, Japan; Department of Dentistry, University of California San Francisco, CA, USA; Departments of Medicine and Endocrinology, University of California San Francisco and Veterans Affairs Health Center San Francisco, CA, USA

## Abstract

Postnatal cell fate has been postulated to be primarily determined by the local tissue microenvironment. Here, we found that Mediator 1 (*Med1*) dependent epigenetic mechanisms dictate tissue-specific lineage commitment and progression of dental epithelia. Deletion of *Med1*, a key component of the Mediator complex linking enhancer activities to gene transcription, provokes a tissue extrinsic lineage shift, causing hair generation in the dental environment. *Med1* deficiency gives rise to unusual hair growth via primitive cellular aggregates on incisors. Mechanistically, we found that Med1 establishes super-enhancers that control enamel lineage transcription factors in dental stem cells and their progenies. However, *Med1* deficiency reshapes the enhancer landscapes and causes a switch from the dental epithelial transcriptional program towards hair and epidermis on incisors *in vivo*, and in dental epithelial stem cells *in vitro. Med1* loss also provokes an increase in the number and size of enhancers. Interestingly, control dental epithelia already exhibit enhancers for hair and epidermal key transcription factors; these expand in size and transform into active super-enhancers upon *Med1* loss suggesting that these epigenetic mechanisms cause the transcriptomic and phenotypic shift towards epidermal and hair lineages. Thus, we propose a role for Med1 in safeguarding lineage specific enhancers, highlight the central role of enhancer accessibility and usage in lineage reprogramming and provide new insights into ectodermal regeneration.

## INTRODUCTION

Postnatal cell fates are controlled by selective transcriptional programs, that are coordinated by an underlying epigenetic machinery and cellular microenvironments^1-4^. Ectoderm evolves into multiple lineages including enamel forming dental lineages, or to hair and epidermis generating skin epithelia^5-7^. Adult stem cells are endowed with tissue specific potential maintaining however the ability to differentiate into different cell types^8-10^To date, the mechanisms controlling specific cell lineage are yet not fully understood, and accordingly, the factors safeguarding dental or epidermal cell lineages are still largely elusive. Understanding these mechanisms is also crucial for our efforts in combating diseases like alopecia or tooth loss as it might create new entry points for the development of bioengineering strategies for the regeneration of tissues like tooth enamel and skin hair, which have been challenging^7^.

The mouse incisor provides an excellent model system to study postnatal lineage commitment and progression. Dental epithelial stem cells (DE-SCs) residing in the cervical loop (CL) at the proximal end of incisors support the continuous growth of the incisors throughout the lifespan of a mouse. They regenerate the enamel organ by giving rise to all the dental epithelia such as enamel matrix producing ameloblasts, Notch1 regulated stratum intermedium (SI) and others^11-16^.

Skin epithelia derived from the same ectoderm are divided into the interfollicular epidermis and hair follicles during embryonic stages of skin formation^6^. Postnatally, hair follicle stem cells residing in the bulge region of hair follicles regenerate hair in skin^17,18^. Hair follicles are essential for hair growth and regeneration in mammalian skin and are characterized by a complex multilayered bulb structure containing the outer and inner root sheath, medulla, and cortex^19,20^. Following the initial developmental cycle, hair is maintained by hair follicle cycling in mice, characterized by an anagen phase during which the hair follicle expands, followed by a transitional catagen phase, leading to the involution of the hair follicle, and telogen, the resting phase. The cycle then begins a new in response to signals from the dermal papilla, a specialized mesenchymal group of cells attached to the proximal end of the hair bulb^21,22^. The shift between these phases involves substantial changes in gene expression patterns, which can be activated by injury. For example, hair depilation induces new hair growth by stimulating anagen with induction of gene expression such as those encoding hair keratins^23^.

Several transcription factors and pathways have been shown to induce dental as well as hair and epidermal cell differentiation. During embryonic development, dental and skin epithelia share similar transcriptional networks as they are derived from the same ectoderm. Postnatally, each lineage is controlled by specific transcription factors. For example, paired like homeodomain 2 (*Pitx2)*^*24*^ and Nk homeobox 3 (*Nkx2-3)*^*25*^ ISL LIM homeobox 1 (*Isl1)*^*26*^ are implicated in enamel epithelial homeostasis^7^ while transcription factors and signaling genes including HR lysine demethylase and nuclear receptor corepressor (*Hr also called hairless*) and Ras and Rab interactor 2 (*Rin2*) are important in controlling hair differentiation because their mutation causes alopecia (hair loss)^27,28^. On the other hand, AP1 factors (*Fos/Jun*) and the tumor proteins of the p53 family including *Tp63* are crucial to direct postnatal epidermal differentiation in the skin^20^. However, the check point factors specifying dental vs epidermal cell fate are still largely elusive.

Mediator is a promising candidate to specify the ectodermal lineages. Mediator complex controls cell lineages by facilitating the expression of cell fate driving transcription factors and genes through large cluster of enhancers called super-enhancers, that differs from typical enhancers in size and an ability to induce transcription of cell identity genes^4,29^. Mediator 1 (Med1) is a subunit of the Mediator complex, which facilitates gene transcription by linking enhancers to the RNA polymerase complex at transcriptional start sites (TSS) in gene promoters. Med1 has been shown to localize at super-enhancers and typical enhancers which represent distal regulatory elements recognized as key epigenetic hubs^4,29^. In fact, Med1 assures pluripotency of embryonic stem (ES) cells by enforcing the expression of Klf4, Oct4, Sox2, and Nanog, four key reprogramming factors. In addition, reduced expression of Mediator subunit like *Med1* triggers differentiation of ES cells^4,29^. Deletion of *Med1* also affects somatic cell fates such as epidermal cells in skin^30-34^. *Med1* ablation hinders invariant natural killer T cell development^35^ and prevents luminal cell maturation in the mammary epithelia in mice^36,37^. These data indicate a role for *Med1* in controlling lineage commitment in somatic cells as well. However, the exact role of *Med1* is not fully understood as previous biochemical studies indicated that removal of Med1 disturbs selective transcription^38^, but is dispensable for Mediator complex formation^38^.

In the present study, we investigate the role of *Med1* in governing ectoderm lineages using a conditional knockout (cKO) mouse model, in which *Med1* expression is deleted from ectoderm derived epithelial cells. *Med1* ablation causes hair generation in incisors with loss of the ability of dental epithelia to produce enamel resulting in severe enamel dysplasia^33,39,40^. Here, we present a unique cellular process by which *Med1* ablation causes ectopic hair growth on incisors. Using an integrative multi-omics approach, we also demonstrate that this intriguing phenomenon is due to a reshape of the enhancer landscape in which ectoderm conserved enhancers are amplified to induce epidermal and hair driving transcription factors.

## RESULTS

### Lack of *Med1* in dental epithelia causes ectopic hair without hair follicles on incisors

During postnatal murine tooth maintenance, CL residing dental epithelial stem cells (DE-SCs) differentiate into Notch1 regulated SI, ameloblasts and other epithelia to mineralize enamel on the incisor surface (Fig. 1a, top). Intriguingly, deletion of *Med1* from *Krt14* expressing DE-SCs and their progenies provoked a major phenotypical shift in the dental compartment causing enamel formation to be replaced by unusual, ectopic hair growth on mouse incisors (Fig. 1b)^33^. As hair growth on the skin depends on multiple cell populations arranged to form hair follicles (Fig. 1a, bottom), the question arises on how hair development is achieved in a dental environment. Therefore, we evaluated the cellular process leading to hair development in *Med1* cKO incisors by comparing it with physiological hair in skin. Although hair grown on incisors of our mouse model showed a comparable hair architecture (cuticle), composition (guard hair and zig-zag hair) and morphology as naturally grown skin hair (Fig. 1b), hair formation significantly differed on incisors in which hair supporting tissues and root structures lacked hair follicles (Fig. 1b black follicles). In fact, while skin grown hair is regenerated from large anagen follicles in response to signals from the dermal papilla and maintained by small telogen follicles (Fig. 1c, bottom orange arrows), we found that dental hair is supported by atypical cell clusters without typical hair follicle-like structures in *Med1* cKO incisors at 4wk when hairs were first visible (Fig. 1c Top, blue arrows with dotted lined region, and Suppl. Fig. 1e). These abnormal cell clusters originated from disorganized SI layer (Suppl. Fig. 1b red triangles) that progressed into expanded papillary layers (Suppl. Fig. 1c yellow triangles) to form the hair bases in dental mesenchymal tissues (Suppl. Fig. 1e yellow triangles). NOTCH1 positive SI/SR derived papillary cells formed unusual cell aggregates and differentiated into hair lineage as they express the hair marker keratin 71 (KRT71) at 4wk at maturation stage (Suppl. Fig. 1d). At the adult stages (3 months of age and later), these aggregates formed aberrant pocket-like structures (Fig. 2a). These NOTCH1 expressing SI/SR derived cell clusters gave rise to hair KRT71 expressing cells as well as in cells expressing the epidermal marker Loricrin (LOR) at 3 months of age (Fig. 2a, b). However, the spatial organization of these cells was scattered and randomly distributed (Fig. 2b two left panels, large blue arrows), that differed from the well-defined cellular framework found along hair follicles and on the surface of the interfollicular epidermis in the skin (Fig. 2b, right panels, orange arrows). Furthermore, hair depilation of skin initiated the hair cycle by enlarging hair follicles (anagen) (Fig. 2c left 3 panels, orange arrows) and inducing the expression of *Krt31* (Suppl. Fig. 2a skin). In contrast, dental hair depilation did not activate any follicle-like structures in the hair supporting tissues on *Med1* null incisors (Fig. 2c two right panels, blue arrows), in either distal or proximal incisor regions (Suppl. Fig. 2b) nor is the expression of the hair marker *Krt31* induced (Suppl. Fig. 2a tooth). Nevertheless, hair grew back at its original length 12 days after depilation in *Med1* cKO incisors (Fig. 2d blue arrow), as occurred in normal skin (Suppl. Fig. 2c).

**Fig. 1:**
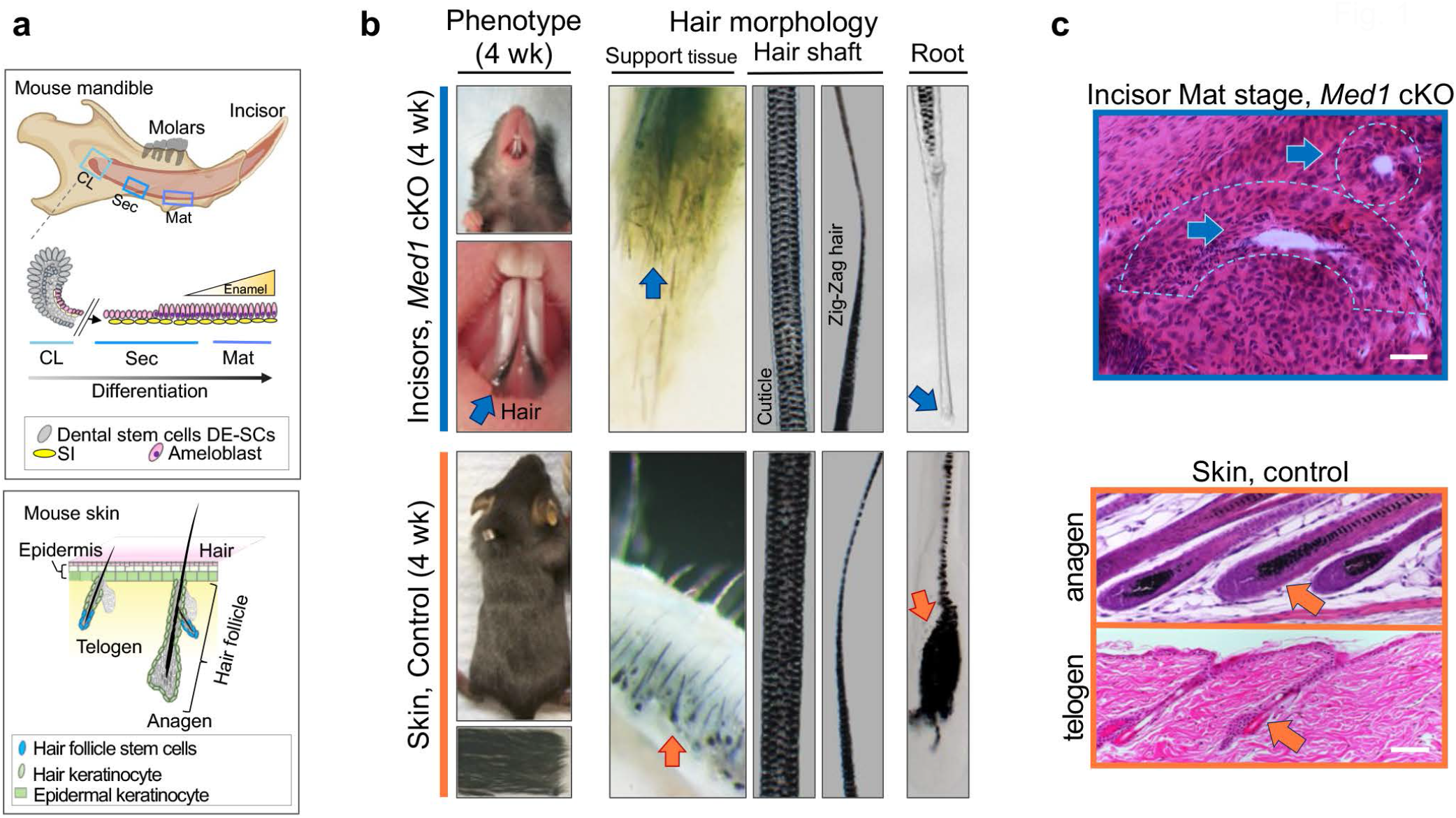
Loss of *Med1* in dental stem cells causes ectopic hair growth on incisors. **a** Top, diagram representing normal dental epithelial differentiation and enamel formation in mouse mandible. CL, cervical loop; Sec; secretory stage; Mat, maturation stage; DE-SCs, dental epithelial stem cells; SI, stratum intermedium. Bottom, hair regeneration by hair follicles under cycling regulation of anagen and telogen in skin. **b** Hair growth on *Med1* cKO incisors at 4 weeks of age (top) is compared to normal hair in skin (bottom). **c** Top, H&E sections of tissues supporting hair growth (blue arrows with dotted lines) in *Med1* cKO incisors at 4 weeks of age. Bottom, H&E sections for hair follicles at anagen and telogen phases supporting hair growth and regeneration in the skin (orange arrows). Bars = 25 μm. For (**b, c**), representative images are shown.

**Fig. 2:**
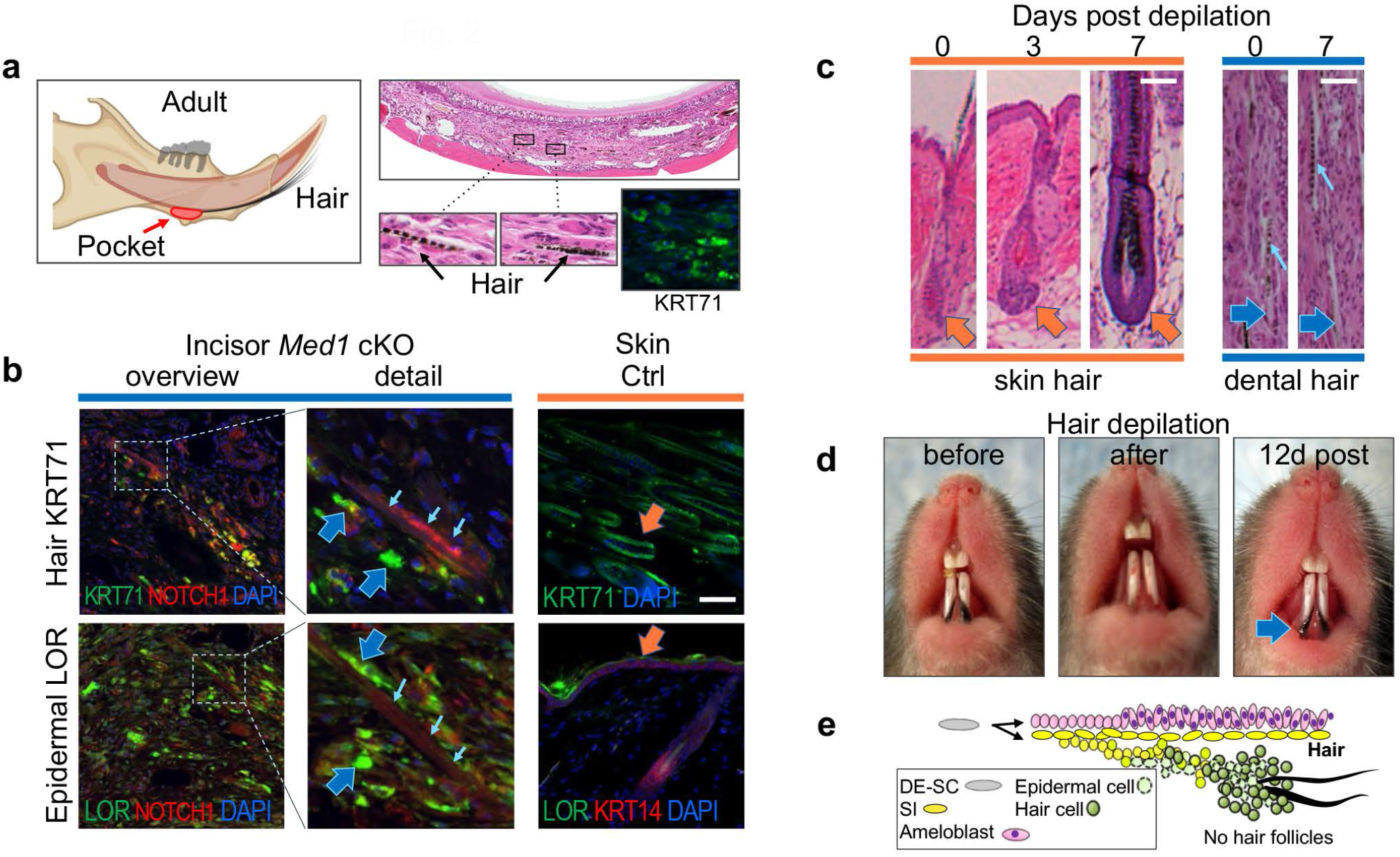
*Med1* null incisors develop and regenerate hair without hair follicles. **a** Left, diagram depicting hair generation via pocket-like structures (red) in *Med1* cKO incisors in adult mice. Right, H&E staining to evidence hair in dental tissues and immuno-staining against hair marker KRT71 (green). **b** Top, immuno-staining of hair marker KRT71 (green) and NOTCH1 (red) in the aberrant pockets on *Med1* cKO incisors or of KRT71 only (green) in normal skin. Bottom, epidermal marker Lor (green) and NOTCH1 (red) in dental tissues compared to Lor (green) and KRT14 (red) localization the skin. Small blue arrows show auto-fluorescent hair structures. **c** H&E staining sections of skin (orange) and dental (blue) sections before (day 0) and after hair depilation (3 and 7 days). Small arrows show hair. Bars = 25 μm. **d** Full hair regeneration 12 days after hair depilation from *Med1* cKO mice (blue arrow). **e** Schematic representation of cellular processes and anatomical location of dental SI/SR derived dental epithelia that are gradually transformed into epidermal (pale green) and hair gene expressing (green) aberrant cell aggregates that lack hair follicle structures in *Med1* cKO incisors. For **a**-**d**, representative images are shown, and reproducibility was confirmed at least in two different litters of *Med1* cKO and control mice.

These results demonstrate that hair can grow and is regenerated through SI derived dental epithelia even in the absence of well-organized hair follicle structural features (Fig. 2e). Expression of hair and epidermal markers in the hair generating cell aggregates further indicates that the presence of certain transcriptional patterns allow for the generation of hair via a primitive cellular environment.

### *Med1* deficiency is sufficient to switch the dental transcriptional program towards hair and epidermis in incisors *in vivo*

Cell identity and cell lineage progression tightly associate with a timed expression of specific gene-sets. As loss of *Med1* causes hair growth in the dental compartment, using total gene expression datasets we investigated the expression patterns of epidermal and hair related gene sets at distinct anatomical locations of mouse incisors in *Med1* cKO incisor (Fig. 3a, diagram). Our analyses revealed that compared to control mice, in *Med1* cKO mice some epidermal genes are already induced in the DE-SCs containing CL region at 4 weeks of age (Fig. 3b left panel, blue dotted box, Fig. 3c pale blue bar). Subsequently, as DE-SCs differentiate towards the secretory (Sec) and maturation (Mat) stages, expression of many epidermal genes strongly increased in *Med1* deficient tissue (Fig. 3b). Intriguingly, hair related genes were not yet detected in the CL locus but become clearly upregulated downstream in the Sec and Mat regions (Fig. 3b right panel, Fig. 3C blue bar). Immunostaining confirmed the sequential induction of epidermal and hair markers as the epidermal markers LOR and KRT1 were detected at secretory stage^39^ but KRT71 was expressed only at maturation stage (Suppl, Fig. 1d). These data illustrate how in cKO mice the expression of selected gene-sets during dental hair development parallels embryonic and postnatal skin development in wild type mice where epidermal development occurs first (E15) (Fig. 3c red bar), followed by hair follicle formation (E18) (Fig. 3c orange bar). These results demonstrate that *Med1* deficiency is sufficient to implement epidermal and hair related transcriptional programs priming dental stem cells towards skin epithelial differentiation *in vivo*.

**Fig. 3:**
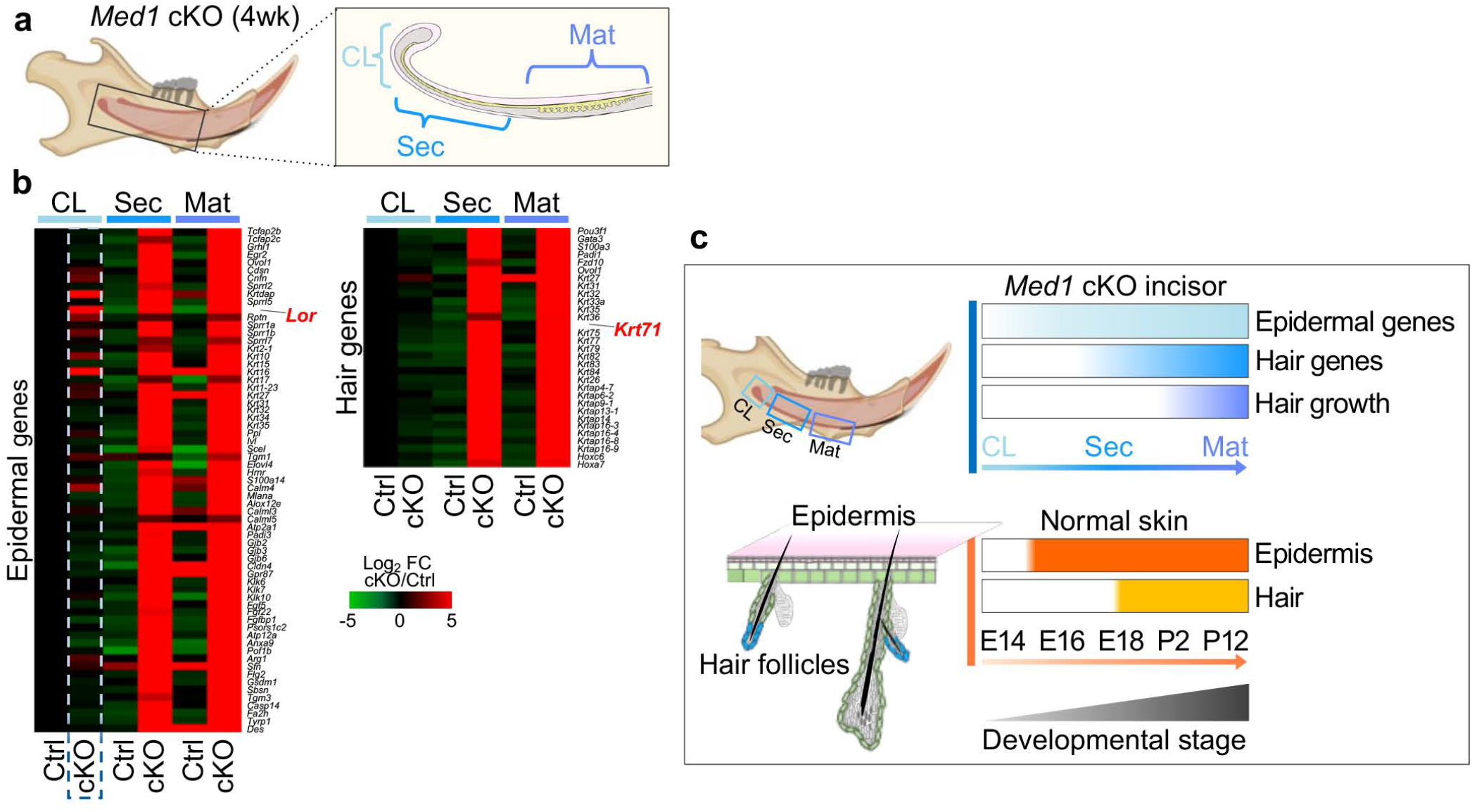
Loss of *Med1* activates epidermal and hair gene expression in developing incisors. **a** Diagram depicting the anatomical locations tested in Ctrl and *Med1* cKO incisors. CL, cervical loop; Sec, secretory stage; Mat, maturation stage. **b** Heatmap showing differential gene expressions for epidermal genes (left) and hair genes (right) in *Med1* cKO incisors versus Ctrls at 4 weeks of age. For each gene, fold changes are compared to CL Ctrl; n=3 for all samples and average values for each group are shown. **c** Top, diagram depicts sequential expression of epidermal genes (light blue) and hair genes (blue) as well as hair growth (purple) in *Med1* cKO mice in regard to the different anatomical locations on incisors (CL, Sec and Mat) of *Med1* cKO incisors. Bottom, pattern comparison to epidermal (orange, E15) and hair gene inductions (yellow, E18) during embryonic development of the skin.

### *Med1* deletion redirects dental epithelial stem cells towards an epidermal fate in *vitro*

Although deletion of *Med1* causes a lineage shift *in vivo*, the question arises if aberrant dental hair growth is due to lineage predisposition of dental epithelial stem cells. To investigate this phenomenon, dental stem cells from CL tissues were cultured. CL tissues from 8-10 weeks old control and *Med1* cKO mice were micro-dissected and digested cells were plated until self-renewing stem cell colonies formed (Fig. 4a, b). An enhanced proliferation rate was observed for the *Med1* deficient cells as shown by the increased number of BrdU positive cells as well as by the generation of larger colonies (Fig. 4b, cKO). Confirming and expanding these results, IPA (ingenuity pathway analysis) of whole transcriptomic data from these cultures not only revealed an upregulation of cell growth and proliferation related pathways in *Med1* null cells, but also showed that these cells spontaneously differentiate into the epidermal lineage in monolayer culture (Fig. 4c, d), even without factors or mesenchymal feeder cells to induce epidermal differentiation. In fact, like Sec tissue from cKO mice (Fig 4d middle column), an extensive set of epidermal related genes are clearly upregulated in cultured *Med1* null DE-SCs as well (Fig. 4d right column). Of note, in contrast to our *in vivo* data shown above, genes related to hair differentiation are only partially induced *in vitro* (Fig. 4d), suggesting that full hair differentiation requires additional external processes. Furthermore, IPA analysis also identified members of the p53 family (Tp63/53/73) as top upstream inducers of epidermal differentiation (Fig. 4e) that are linked to the key epidermal AP-1 transcription factors *Fos* and *Jun* (Fig. 4f), resembling their roles in the skin. Collectively, these results show that Med1 deletion reprograms DE-SCs towards epidermal fates through a cell intrinsic mechanism. Together with our *in vivo* observation in *Med1* cKO, these results lead us to investigate the underlying epigenetic and transcriptional mechanism, by which Med1 assures enamel lineage and Med1 deficiency causes a lineage shift.

**Fig. 4:**
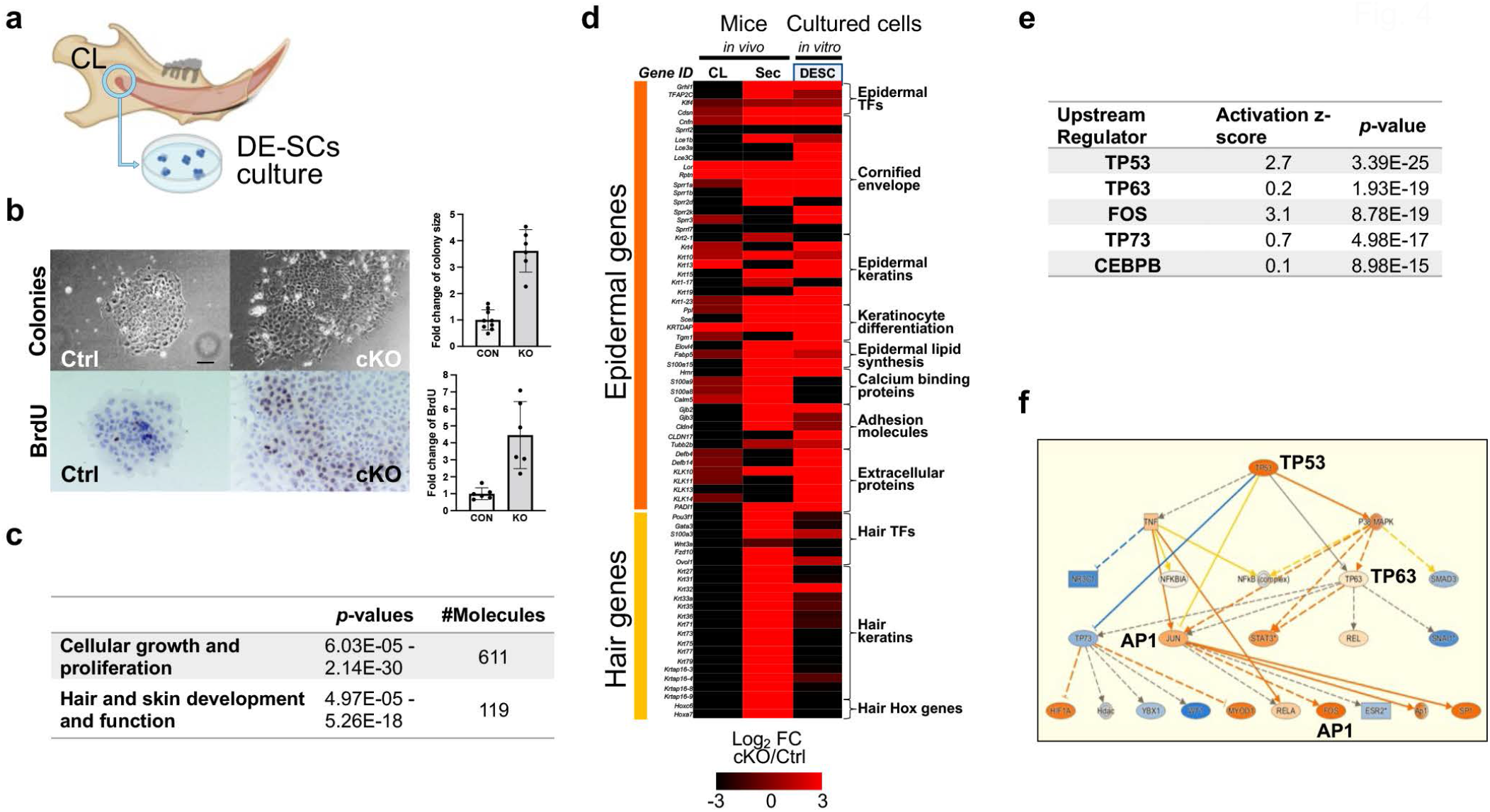
*Med1* deficiency directs DE-SCs towards epidermal fate *in vitro*. **a** Generation of DE-SC culture from CL tissues. **b** Left, representative bright field images of DE-SC colonies and BrdU staining in Ctrl and *Med1* cKO. Bar = 10 μm. Right, quantification of colony size and BrdU positive cells/colony are shown as fold changes (cKO/Ctrl); in both comparisons *p*>0.01. **c** Biological processes associated with loss of *Med1* in DE-SCs as identified by Ingenuity Pathway Analysis (IPA) on microarray gene expression data. **d** Heatmap showing differential gene expressions of epidermal and hair related genes in *Med1* cKO vs Ctrl incisor tissues (CL and Sec *in vivo*) and cultured *Med1* cKO vs Ctrl DE-SCs (third lane); n=3 for all samples and average values for each comparison are shown. **e** Upstream regulators responsible for biological process induced by loss of *Med1* in DE-SCs as identified by IPA on microarray data as used in (**c**). **f** Mechanistic network representation for TP53/63 pathways in induced in *Med1* lacking DE-SC culture regulating epidermal fate driving AP-1 factors; upregulated genes are in orange and direct relationships are shown by solid lines. Reproducibility was confirmed by two independent cultures, and representative data are shown.

### Med1 regulates enamel lineage driving transcription factors through super-enhancers

Lineage commitment and reprogramming are controlled by cell fate driven transcription factors. Mediator including Med1 orchestrates these key factors by associating with super-enhancers and promoters^29^. Therefore, we first identified these key transcription factors by assessing the genome wide distribution of MED1 containing enhancers during dental epithelial development in control mice (4wk). For this purpose, we performed MED1 ChIP-seq of the stem cell containing head region of the CL (CLH) as well as on the CL tail area (CLT) which includes their progenies (Fig. 5a, left diagram). MED1 accumulated in the distal intergenic regions (Suppl. Fig. 3a) where it associated to several hundred super-enhancers in the CLH and CLT tissues (Fig. 5b and Suppl. Fig. 3b). These include super-enhancers near lineage driving transcription factors. In the CLH tissues, these include enamel lineage transcription factors including *Pitx2, Isl1, and Nkx2-3* (Fig. 5b left panel and Fig. 5c left 3 panels), which may prime adult stem cells to an enamel fate, consistent with their essential roles in tooth morphogenesis during embryonic development^25,41^. In contrast, in CLT samples, MED1 super-enhancers were found around key transcription factors like Satb1 homeobox 1 (*Satb1)* and Runx family transcription factor 1 (*Runx1*) and 2 (*Runx2*) which control later cellular processes like differentiation and enamel mineralization^41,42 43^(Fig. 5b right pane and Fig. 5c right 3 panels). Of note, the mRNA expression levels of these transcription factors were strongly reduced in CLT tissues in *Med1* cKO mice (Fig. 5d), further implicating their role in enamel formation as *Med1* null mice show severe enamel dysplasia^40^. In addition, MED1 levels are enriched around the promoters of these transcription factors but not of ameloblast markers (Suppl Fig. 3d), indicating that Med1 directly activates enamel fate related transcription factors rather than supporting late differentiation. These data indicate that Med1 programs dental stem cells and their progenies to commit to and progress the enamel lineage by controlling the expression of lineage driven transcription factors (Fig. 5e).

**Fig. 5:**
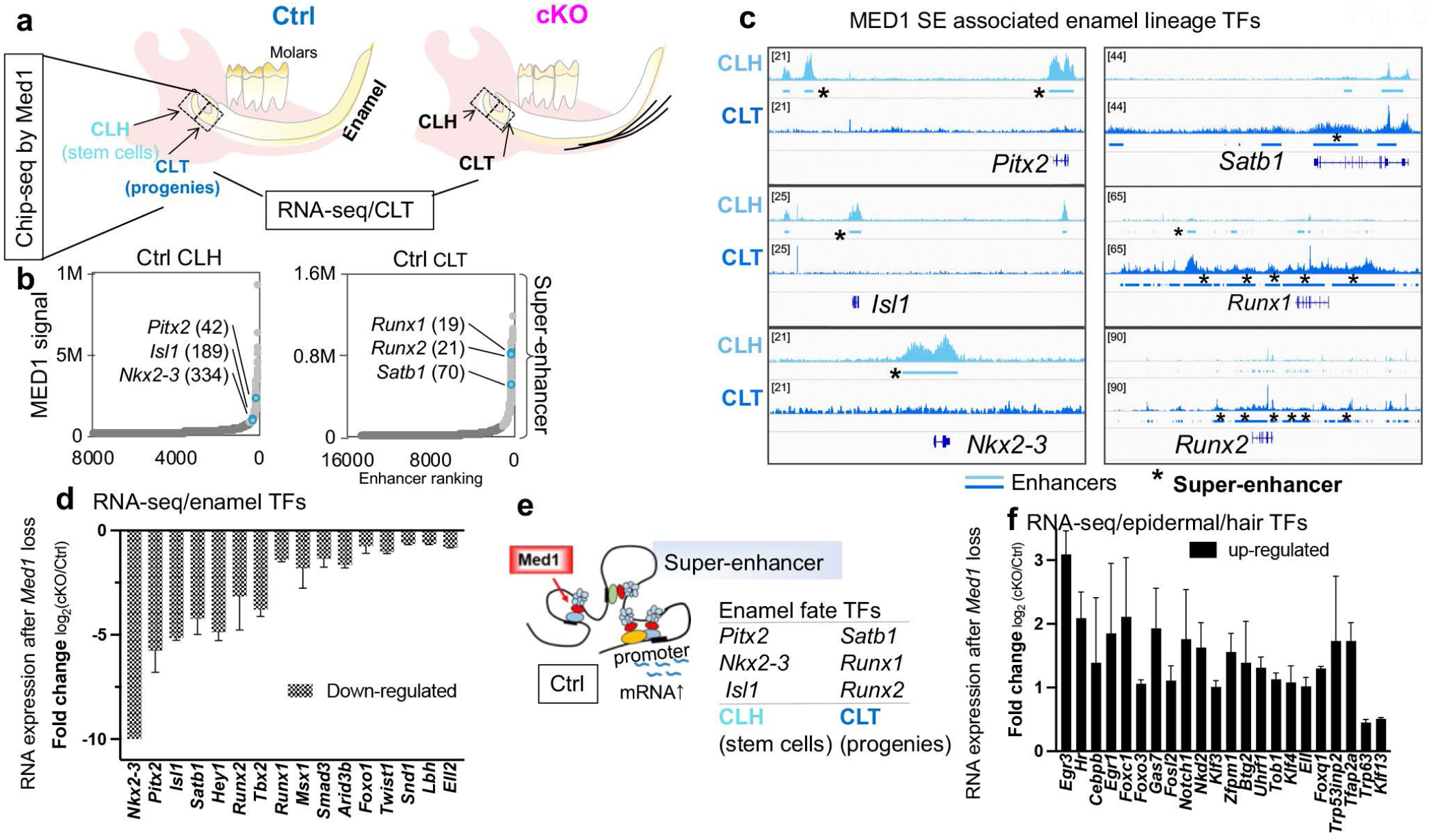
Med1 directly controls enamel lineage transcription factors by associating with their enhancers and promoters. **a** Schematic representation of tissues isolated (CLH stem cells, and CLT their progenies) from normal mouse mandibles for MED1 Chip-seq and RNA-seq. **b** MED1 bound enhancer clustering in CLH and CLT tissues. Light gray and blue dots represent super-enhancers, dark grey dots are typical enhancers; blue dots outline super-enhancer associated enamel transcription factors as designated by gene name and enhancer ranking numbers. **c** Genomic MED1 binding profiles in CLH (light blue) and CLT (blue) tissues for relevant transcription factors. Super-enhancers (SE) are marked by bars with asterisks*. Average profiles from two independent Chip-seq experiments are shown. **d** Transcription factors (TFs) identified through MED1 super-enhancers that were down-regulated in *Med1 cKO* (CLT) as measured by RNA-seq. **e** Schematic representation of Med1 to induce enamel fate transcription factors (TFs); distal Med1 (red) containing super-enhancers associate to gene promoter to induce mRNA expression (blue waves). **f** Enhancer associated epidermal or hair fate transcription factors (TFs) that were upregulated in cKO (CLT) as measured by RNA-seq. **d, f** Data are shown as average and SD of fold changes (log_2_FC) in cKO compared to Ctrl within two different litters of *Med1* cKO and control mice with statistical significance (*p*<0.05).

Intriguingly, we also found enhancers around a different set of transcription factors (Suppl. Fig. 3c pink labeled) involved in epidermal differentiation of the skin including the genes of early growth response (*Egr3)*, CCAAT enhancer binding protein β (*Cebpb*), Fos like 2 (*Fosl2*) (AP-1 transcription factor subunit), Kruppel like factor 3 (*Klf3*), Forkhead box C1 (*Foxc1*), *Foxo3*, tumor protein *Tp63*, and hair fate driving factors like *Hr* and *Rin2*. Their mRNA expressions were substantially upregulated in *Med1* cKO mice at CLT tissues (Fig. 5f), implicating their roles to induce epidermal and hair fates in *Med1* cKO mice.

### Med1 deficiency reshapes the enhancer landscape in DE-SCs and their progenies

As loss of *Med1* inhibits the expression of Med1 associated enamel lineage driving transcription factors while inducing epidermal transcription factors in CL tissues, we next probed if this is linked to epigenetic changes in promoter and enhancer patterns upon loss of *Med1*. To test this hypothesis, we performed ChIP-Seq analysis against an alternative enhancer marker, histone 3 lysine 27 acetylation (H3K27ac), in CLH and CLT tissues of 4-wk-old *Med1* cKO mice and their littermate controls (Fig. 6a) as the Med1 antibody is not useful for *Med1* cKO tissues. It is known that histone acetylation (H3K27ac) generally co-localizes with Mediator such as Med1 at distal regulatory elements like typical and super-enhancers and acts as a major inducer of gene expression by increasing chromatin accessibility^29,44^. Compared to controls, we found major genomic changes in the H3K27ac binding patterns of *Med1* cKO CLT tissues. These were associated with biological functions such as enamel mineralization and tooth morphogenesis (Fig. 6b blue bars), as well as epidermal and hair development (yellow bars), consistent with the observed *Med1* cKO phenotypes. These results suggest that phenotypic and transcriptional changes are due to epigenetic changes. In addition, *Med1* deletion substantially increases the number of super-enhancers (Fig. 6c) as well as typical enhancers (Suppl Fig. 4a) in both CLH and CLT. Some of these new enhancers are found around loci coding for epidermal and hair lineage genes (Fig. 6c bar graphs) that become upregulated upon loss of *Med1* in CLT tissue. For example, a new super-enhancer was formed in the *Med1* cKO on epidermal transcription factor *Fosl2* (Ap-1 factor) loci (Fig. 6d blue bar in boxed region), directly linking its increased mRNA expression to the loss of *Med1* (see Fig. 5f and Suppl Fig. 4b). Corroborating these results and in line with our transcriptional network studies in *Med1* null cell cultures (see Fig. 4f), we also found that the binding motifs for the epidermal inducible transcription factors, Tp53/63, Egr1 and AP-1 are significantly enriched in super-enhancers in cKO at CLT compared to controls (Fig. 6e). Collectively, these data demonstrate that loss of *Med1* alters the epigenetic landscapes in DE-SCs and their progenies. These results prompted us to investigate the epigenetic mechanism by which Med1 deficiency induces epidermal and hair lineages more in detail.

**Fig. 6:**
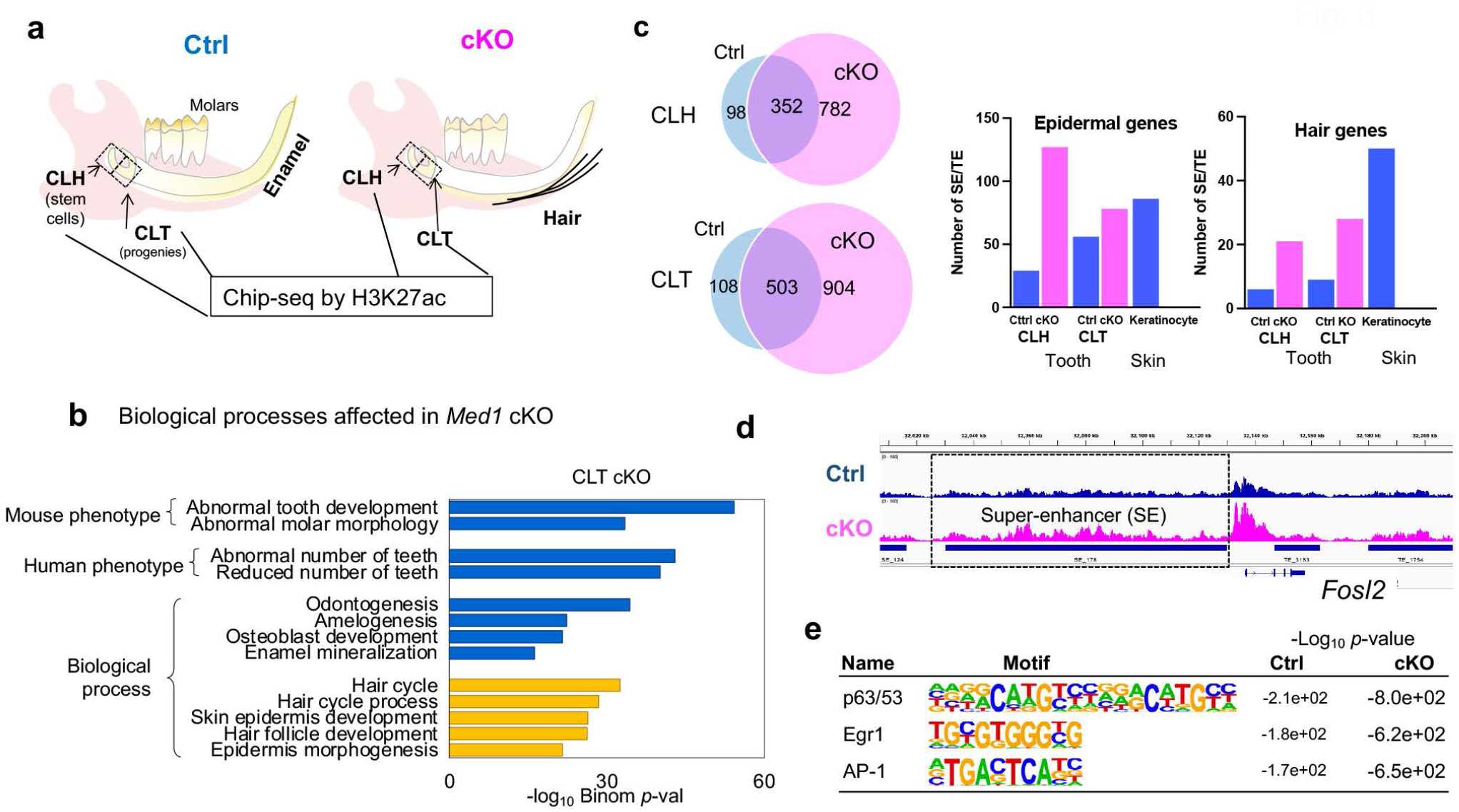
Loss of *Med1* expands the super-enhancer landscape in CLH and CLT tissues. **a** Schematic representation for the isolation of CLH (stem cells) and CLT (their progenies) from *Med1* cKO and littermate Ctrl mandibles for Chip-seq against H3K27ac to compare actively transcribed genomic regions. **b** GREAT based GO analysis for genes associated with differential H3K27ac peaks to elucidate biological processes affected by *Med1* loss in CLT tissues. **c** Number of super-enhancers in Ctrl (blue) and cKO (pink) tissues. Bar charts comparing typical and super-enhancers associated with epidermal and hair related genes before and after loss of *Med1* in CLH and CLT tissues. As a comparison, data from skin derived keratinocytes are included. **d** Chip-seq profiles for H3K27ac occupancy for *Fosl2* gene, in which the super-enhancer in cKO is marked by blue line. **e** Most enriched transcription factor binding motifs found in super-enhancers formed in *Med1* cKO compared to Ctrl in CLT tissues; statistical significance are shown as log_10_(*p*-values). All the Chip-seq data are average of duplicates conducted in 2 different litters of *Med1* cKO and littermate controls.

### *Med1* deficiency induces epidermal and hair driving transcription factors via amplification of ectoderm conserved enhancers

During embryonic development, dental and skin epithelia are derived from the same ectoderm by sharing transcriptional networks. Our results indicated that normal dental epithelia already possess enhancers around key epidermal and hair transcription factors (Suppl Fig. 3c pink labeled). We further found that even though dental epithelia usually do not commit to epidermal fates, they widely share the enhancers with skin epithelia both epidermis and hair follicle derived cells^45^ for most of the transcription factors induced upon *Med1* loss (Fig. 7a). Importantly, *Med1* deficiency elevated and upgraded these shared enhancers into transcriptionally active super-enhancers as outlined for the *Hr* (hairless) locus (Fig. 7b pink bar and Suppl Fig. 4c). Loss of *Med1* also expanded enhancers in size for epidermal factors *Cebpb* and *Foxo3* loci (Suppl. Fig 4d). In addition, enhancers near many hair-lineage related genes were elevated to super-enhancers upon the loss of *Med1* in CLT (Fig. 7c and Suppl. Fig. 4e) which coincided with an induction at their mRNA levels that were measured by RNA-seq (Fig. 7d and Suppl. Fig. 5a). Intriguingly, these factors controlling hair lineage were shown to be regulated by super-enhancers in the skin before^4^. Besides the generation of new super-enhancers, we also found an increase in H3K27ac levels around promoters of these genes (Suppl. Fig. 4f), which further positively corelated with an increased gene expression (Fig. 7e). In *Med1* cKO CLH tissues, increased H3K27ac promoter levels positively correlated with an elevated expression of genes involved in preventing ossification (Suppl. Fig. 5a, and 6b, c), and H3K27ac occupancy increased in ossification inhibiting transcription factors of *Sox9* and *Tob1* loci in their enhancers (Suppl. Fig. 6c). This further corroborates the dental phenotype of our mouse model which is not only characterized by hair growth but also by enamel dysplasia. These results demonstrated that cell fate switch is at least in part due to an inflation of developmentally conserved epidermal and hair enhancers.

**Fig. 7:**
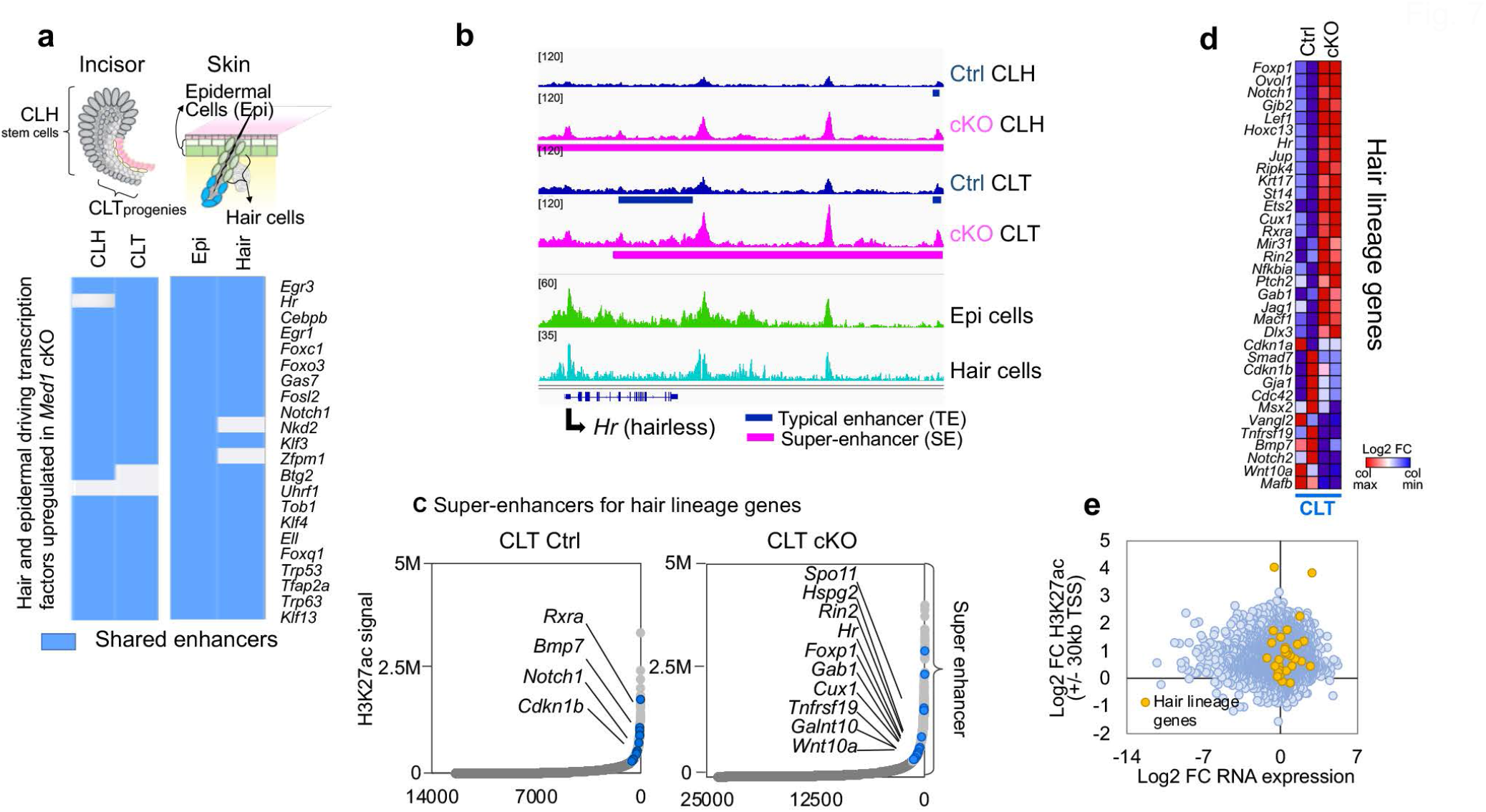
Loss of *Med1* advances pre-existing enhancers to super-enhancers near epidermal and hair lineage genes. **a** Top, Schematic representations of cell sources of dental and skin epithelia are shown. Bottom, heatmap showing shared enhancers between dental and skin epithelia for epidermal and hair lineage related transcription factors that are upregulated in *Med1* cKO. Shared enhancers in dental (CLH, CLT), and skin epithelia including epidermal keratinocytes (Epi), and transient amplifying (TAC) hair follicle keratinocytes (hair) are aligned. **b** Chip-seq profiles around the *Hr* (hairless) locus in CLH and CLT from *Med1* cKO (pink) and Ctrl (blue) mice, compared to epidermal cells (green) and hair TAC keratinocytes (light blue) from skin. Pre-existing enhancers in Ctrl (blue bar) are develop into super-enhancers (pink bar) upon loss of *Med1* cKO in CLT. **c** Enhancer distribution profiles; super-enhancers associated hair lineage genes and exclusively found in Ctrl or *Med1* cKO CLT tissues are noted by the name of neighboring gene (blue circles). **d** Heatmap depicting differential gene expression of hair lineage genes in *Med1* cKO compared to Ctrl in CLT. **e** Correlation between gene expression and H3K27ac promoter occupancy (TSS +/-30 kb) in *Med1* cKO vs Ctrl in CLT tissues for the hair lineage driving gene set (orange dots) compared to all the other genes (blue dots). FC, fold change.

In summary, we propose an epigenetic model in which *Med1* safeguards enamel lineage commitment and progression of dental stem cells and their progenies. *Med1* deletion shifts dental stem cells to epidermal fates by amplifying the ectodermal conserved enhancers around hair and epidermal genes.

## DISCUSSION

Adult stem cell fate is heavily linked to their tissue of origin^2,3,46^. It is postulated that the local cellular and endocrine microenvironment primes and directs tissue specific lineage commitment and differentiation via cellular, transcription and epigenetic mechanisms^2,4^. Here we show that deletion of *Med1*, a key component of enhancer associated Mediator complex, causes ectopic hair growth on incisors. Unlike hair growth in the skin which relies on functionally structured follicles, dental hair is generated from primitive, cellular aggregates without hair follicle morphogenesis. Furthermore, as shown by our depilation experiments, dental hair regrows without induction of the hair cycle, a follicle dependent process required for regeneration of hair in the skin. Thus, our results question the current knowledge that mammalian hair can only be generated from hair follicles consisting of multiple cell layers bearing defined functions. In addition, our study also illustrates how deletion of *Med1* induces lineage shift *in vivo*. Despite cellular agglomerates causing hair growth on incisors exhibit a scattered cellular setup, the mRNA expression profiles found in these structures mimic those of dedicated hair and epidermal cells found in skin epithelia. Thus, while *Med1* negative DE-SCs reside in the dental environment, they are driven towards lineages characteristic for skin epithelia.

Lineage commitment and differentiation are orchestrated by epigenetic programs and regulatory elements including transcription factors and enhancers^4,29,44^. Our results elucidate that already at early dental differentiation stages MED1 containing super-enhancers selectively populate genomic loci coding for enamel fate driven transcription factors in DE-SCs and their progenies. This outlines how *Med1* controls and programs enamel epithelia’s differentiation at multiple stages as it primes DE-SCs towards the dental lineage via activation of *Nkx2-3, Pitx2* and *Isl1* and calibrates lineage progression of DE-SC progenies by regulating the expression of *Runx1/2* and *Satb1*. These findings are consistent with previous studies that *Nkx2-3* controls cusp formation in embryonic dental epithelia^25^, and that *Nkx2-3, Pitx2 and Runx2* are essential for dental development^24,25,41^. *Isl1* is required for patterning and mineralization of enamel^26^, Runx2 for enamel mineralization ^41,42^, and *Satb1* for cell polarity and enamel matrix secretion in amelogenesis^43^.

Intriguingly, although MED1 is a well-recognized super-enhancer component, loss of *Med1* in CL tissues does not compromise super-enhancer formation which is consistent with previous observation^38^, but significantly widens the genomic enhancer landscape instead. In the dental setting, this does not increase the potency of DE-SCs but rather drives these cells into epidermal linages. This coincides with a shift in the super-enhancer profiles around loci coding for transcription factors like *Hr* (hairless) and *Rin2* which are essential for hair formation^27,28^ as well as *Egr1/3, CEBP/a, CEBP/b*, and *Fosl2* which control epidermal differentiation in skin^47,48^. Furthermore, our data show that Tp63/53 binding sites are enriched in super-enhancers in *Med1* cKO mice. These are master regulators for the development of stratified epithelia *in vivo*^20^ and drive epidermal differentiation *in vitro*. In fact, *Med1* null mice activate Tp63 driven epidermal differentiation in the skin, resulting in an accumulation of epidermal markers and lipids in upper hair follicle keratinocytes resulting in hair loss^33^.

Building on our previous studies^39^, these results elucidate a central role for Med1 in assuring enamel lineage. Our results also reveal that basal enhancers are already present at hair and epidermis driving transcription factor loci in control dental epithelia suggesting that a basal epigenetic memory granting multipotency is already established in pre-occurring ectoderm during embryonic development. Consequently, Med1 guarantees adult stem cells to commit to enamel lineage by restricting super-enhancer formation to lineage specific loci. In addition, as loss of *Med1* causes DE-SCs to differentiate into a developmentally related lineage might indicate that reprogramming of DE-SCs into lineages originating from the same germ layer (ectoderm) is easier than overcoming more strict and embryonically set epigenetic barriers^29^.

Hair and tooth are difficult organs to regenerate through pluripotent stem cells based regenerative medicine since no clinical trials have been reported even after extensive efforts^49^. A deep understanding of the cellular and epigenetic mechanisms controlling these lineages and their reprogramming creates new entry points for the development of strategies to combat hair and tooth related diseases. Our results establish that hair formation is achievable in tissue outside of the skin, even without the presence of hair follicles. Similarly, hair was also accidentally generated by DE-SCs transplanted into renal capsules during an effort to regenerate enamel^50^. This suggests that hair could be generated via genetically engineered cell clusters or organoids *in vitro* which might be easier than traditional strategies and bypass time consuming re-programming processes. This would be of great benefit as it would greatly simplify *in vitro* hair generation and allow hair growth without hair follicles which are very challenging to regenerate^51^. Similarly, enamel regeneration might also be feasible by using foreskin derived epidermal stem cells. In fact, cultured epidermal stem cells derived from skin have been shown to differentiate into functional ameloblasts *in vitro*^52^. Overall, these findings, elucidate novel possibilities to develop strategies to overcome hair and enamel associated diseases.

## MATERIALS AND METHODS

### Krt14-driven *Med1* cKO mice

Conditional *Med1* knockout (cKO) mice were generated as described previoulsy^33^. Floxed (exon 8-10) *Med1* mice^53^ (C57/BL6 background) were mated with keratin 14 (*Krt14*) promoter driven *Cre* recombinase mice (The Jackson Laboratory, C57/BL6 background). Genotyping was performed by PCR as previously described^33^ and *Cre* negative littermate mice served as controls (Ctrl). All the experiments were approved by the Institutional Animal Care and Ethics Committee at the San Francisco Department of Veterans Affairs Medical Center.

### Dental epithelial stem cell (DE-SC) culture

CL tissue was separated from 10 wk *Med1* cKO and Ctrl mice, and epithelial tissues were separated and treated by dispase (5U/ml Stem Cell Technology) to separate mesenchymal tissues. Epithelial tissues are further dissociated by accutase (Sigma Milipore), and cells were plated under low density to generate colonies and maintained with DMEM media supplemented with recombinant EGF 20 ng/ml (R&D), FGF 25 ng/ml (R&D), 1X B27 supplement (Gibco). After 8 days of culture, number and size of colonies are evaluated, and RNA is prepared for microarray.

### BrdU staining

DE-SC are labeled by BrdU and stained by a BrdU staining kit (Abcam) by following manufacturer’s instruction.

### Chromatin immunoprecipitation

Chromatin IP was performed by using the LowCell# ChIP kit (Diagenode) for H3K27ac, and the iDeal ChIP-kit (Diagenode) for MED1 according to the manufacturer’s protocols with some modifications as described below. Dissected CLH or CLT tissues were cross-linked with 1% paraformaldehyde for 8 min (H3K27ac) or for 15 min (MED1) and subsequently quenched with 0.125M glycine. Epidermal keratinocytes containing stem and their progenies were isolated from mouse skin were fixed with 1% paraformaldehyde for 10 min. Chromatin was then sonicated with a Covaris S2200 ultra-sonicator (Covaris, Inc.) to obtain DNA fragments with an average length of 300±50 bp. The fragment size was verified with an Agilent 2100 Bioanalyzer (Agilent Technologies). Sheared chromatin was thereafter immunoprecipitated with Protein A-coated magnetic beads (Diagenode) preincubated with 4.5 μg/IP MED1 antibody (Bethyl A300-793A) or 3 μg/IP H3K27ac antibody (ab4729, Abcam). Chromatin input samples were used as reference. Complexes were then washed and eluted from the beads with washing buffer and crosslinks were reversed by incubation with proteinase K at 65°C for 4 hours. Immunoprecipitated DNA along with genomic DNA (Inputs) were purified using the IPure v2 kit (Diagenode). IP efficiency was confirmed by qPCR using primers provided in the kits. Experiments were carried out in duplicates. ChIP-Seq libraries were generated using the Accel-NGS 2S Plus DNA library kits (Swift Sciences) and amplified by PCR for 11-13 cycles. High molecular weight smear was removed in the library by right side size selection using SPRI beads (Beckman Coulter). The libraries were quantified by Agilent 2100 Bioanalyzer with the high sensitivity DNA assay and sequenced on an Illumina HiSeq 4000 (UCSF Center of advanced technology) in a single strand 50 bp run.

### Next generation DNA sequence data analysis

For RNA-Seq data analysis, raw reads were assessed for run quality using fastqc version 0.72 followed by alignment to reference genome (mm10) using RNA STAR Galaxy version 2.6.0b-1 with default settings. Differential gene expression analysis was performed using feature Counts Galaxy version 1.6.3 and DESeq2 Galaxy version 2.11.40.2. For ChIP-Seq data analysis, raw reads from MED1 or H3K27ac ChIP-seq sequencing experiments were assessed for quality using fastqc version 0.72, followed by mapping to reference genome (mm10) using BWA Galaxy version 0.7.15.1 with standard settings. Pooled peak calling was performed with MACS2 callpeak version 2.1.0.20140616.0 using broad peak settings and a minimum FDR cutoff of 0.05. Raw-files for available H3K27ac ChIP-Seq datasets in public from wildtype mice hair stem cell progenies were downloaded from the NCBI-SRA database (SRR1573273, SRR1573274) and analyzed as described above. To assess significantly different peak occupancy between two experimental groups, peak files (MACS2) were analyzed with DiffBind Galaxy version 2.6.6.4 or 2.10.0. Peak size and location were visualized with IGV version 2.3.92. MAYO using group auto-scale for comparable visualization of all shown groups.

Peak intensity around TSS (±10 kbp) as well as genomic peak distribution was assessed with CEAS software (Version1.0.0, Cistrome, Liu Lab). Gene-ontology analysis for Med1 or H3K27ac distal intergenic cis-regulatory elements was performed using GREAT version 3.0.0 (Bejerano Lab, Stanford). Super enhancer analysis was performed using the ROSE package and the software NaviSE. Overlapping typical and super-enhancers between the shown comparisons were assessed with Bedtools (version 2.29) using a minimum overlap of 50%. Motif analysis (de novo and known) was performed using HOMER software [32]. Curated gene-sets used for analyses in this study are provided upon requests.

### Reproducibility and statistical analysis

All the experiments using *Med1* cKO and littermate control mice (n=3) were repeated with at least two litters, and reproducibility was confirmed. The experiments using cultured dental epithelia were conducted in duplicates for microarray, and reproducibility was confirmed. Histologic analyses were performed using 2 sections per mouse per each group of cKO and Ctrl, and representative images are shown. Statistical significance was calculated using software integrated methods or two-tailed unpaired Student’s t-test. If not differently noted, differences with a *p-*value of less than 0.05 were considered as statistically significant.

## Data availability

The array data were submitted to a public database (GEO/NCBI/NIH http://www.ncbi.nlm.nih.gov/geo). Data for *Med1* cKO (4wk), are available with accession numbers GSE50672, GSE50671, GSE50670, respectively under the super-series GSE50673. We are pending of RNA-seq and ChIP-seq data submission.

## AUTHOR CONTRIBUTIONS

Conceptualization: Y.O., R.T., K.Y., S.F., P.D., D.D.B; Methodology: Y.O., R.T., T.N., K.Y.; Investigation: Y.O., R.T., T.N., K.Y.; Writing-Original Draft: Y.O. Writing-Review & Editing: R.T., Y.O., S.F., P.D., D.D.B; Funding Acquisition: D.D.B., Y.O., S.F., K.Y., Resources: R.T., K.Y., S.F., D.D.B., Y.O.

## ACKNOWLEDGMENTS

We thank Ms. Sun Hee Kim Wong for in maintaining the mice and technical assistance. We are also grateful to L. Hu, C. Fong for technical support. We also thank Dr A. Van Wijnen for epigenetic supports. This work was supported by the NIH grant R21 DE025357 (YO), R01 AR050023 (DDB), DOD grant CA110338 (DDB), VA Merit I01 BX003814-01 (DDB).

## Supplemental information

### Supplemental methods

#### Histological analysis

Mouse mandibles were dissected from *Med1* cKO and littermate control (Ctrl) mice and fixed in 4% paraformaldehyde at 4**°**C overnight and decalcified in 0.5 M EDTA for two weeks. Subsequently they were dehydrated and embedded in paraffin wax using standard protocols. The whole mandible was then serially sectioned at a thickness of 7 μm (sagittal sections) containing both the CL and enamel organ. Ultimately, slides were stained with hematoxylin and eosin (HE) and imaged with brightfield microscopy.

#### Immunostaining

For immunostaining, paraffin embedded jaw sections were treated with antigen unmasking solutions (Vector lab Citrate-based, H-3300) and subsequently blocked with an avidin/biotin blocking solution (Vector lab, SP-2001) and incubated with primary antibodies against Krt71 (Covance), Notch1 (Cell Signaling) and Loricrin (Covance). They were then developed with the Vectastain Elite ABC kit (Vector Lab) using DAB staining solution (Vector Lab, SK-4100) until brown signals were visible. The sections were counterstained with Gill’s Hematoxylin.

#### Immunofluorescence

Paraffin-embedded mandible sections were pretreated in 10 mmol/L citrate buffer (pH 6.0, Sigma-Aldrich) for 20 min using a microwave for antigen retrieval. The specimens were blocked by Power Block (BioGenex) and incubated with primary antibodies against KRT71 (Progen), KRT1, (Covance), LORICRIN (Covance) and NOTCH1 (Cell signaling). They were subsequently incubated with species specific secondary antibodies conjugated to fluorescent dyes (Invitrogen Molecular probes), including Alexa 594 (red) and Alexa 488 (green) and counterstained with DAPI. Images were taken with a confocal microscope (LSM510, Carl Zeiss).

#### RNA expression analysis

Total RNA was isolated from 4-week-old total CL, CLH, CLT, Sec or Mat tissues of *Med1* cKO and littermate Ctrl mice using the Pico Pure RNA purification kit (ABI). RNA from cultured dental epithelial stem cells was extracted using the RNeasy mini kit (Qiagen). RNAs were analyzed for quality and purity using the Pico Chip kit on an Agilent 2100 Bioanalyzer (Agilent Technologies) to meet the processing criteria of the UCLA Genomic Core facility for the Illumina array platform (Mouse Ref-8 v2.0 Ambion) or RIN score values for RNA-seq. Microarray data generation and primary microarray data analysis was described previously [12]. Microarray data was used for differential pathway analysis with Ingenuity IPA software (Ingenuity) and results were considered significant for *p-*values less than 0.05. Heat maps were then generated with MeV software using fold changes Log2(cKO/Ctrl). For RNA-Seq analysis, RNA integrity was analyzed with a 2100 Bioanalyzer with an RNA 6000 kit (Agilent Technologies). RNA-Seq libraries were prepared using the Illumina TruSeq v2 library preparation kit and sequenced with paired-end 50 bp reads on an Illumina HiSeq 4000 in biological duplicates.

For quantitative real-time polymerase chain reaction (rt-qPCR), cDNA was synthesized from ∼0.5 μg RNA using the SuperScript III first strand synthesis system (Invitrogen) as described by the supplier and subjected to rt-qPCR amplification using the QuantiTect SYBR-Green PCR Kit (Qiagen) and a 7300 or 7500 Real Time PCR system (Applied Biosystems) using the following conditions: 10 min 95°C followed by 45 cycles of 30 sec at 95°C, 30 sec at 60°C and 30 sec at 72°C. Relative mRNA levels were compared to the housekeeping gene *Gapdh* and determined with the ΔΔCt method. Primers for the analyzed genes are provided upon requests. All rt-qPCR assays were performed in triplicates.

## Supplemental Figures

**Suppl. Fig. 1:**
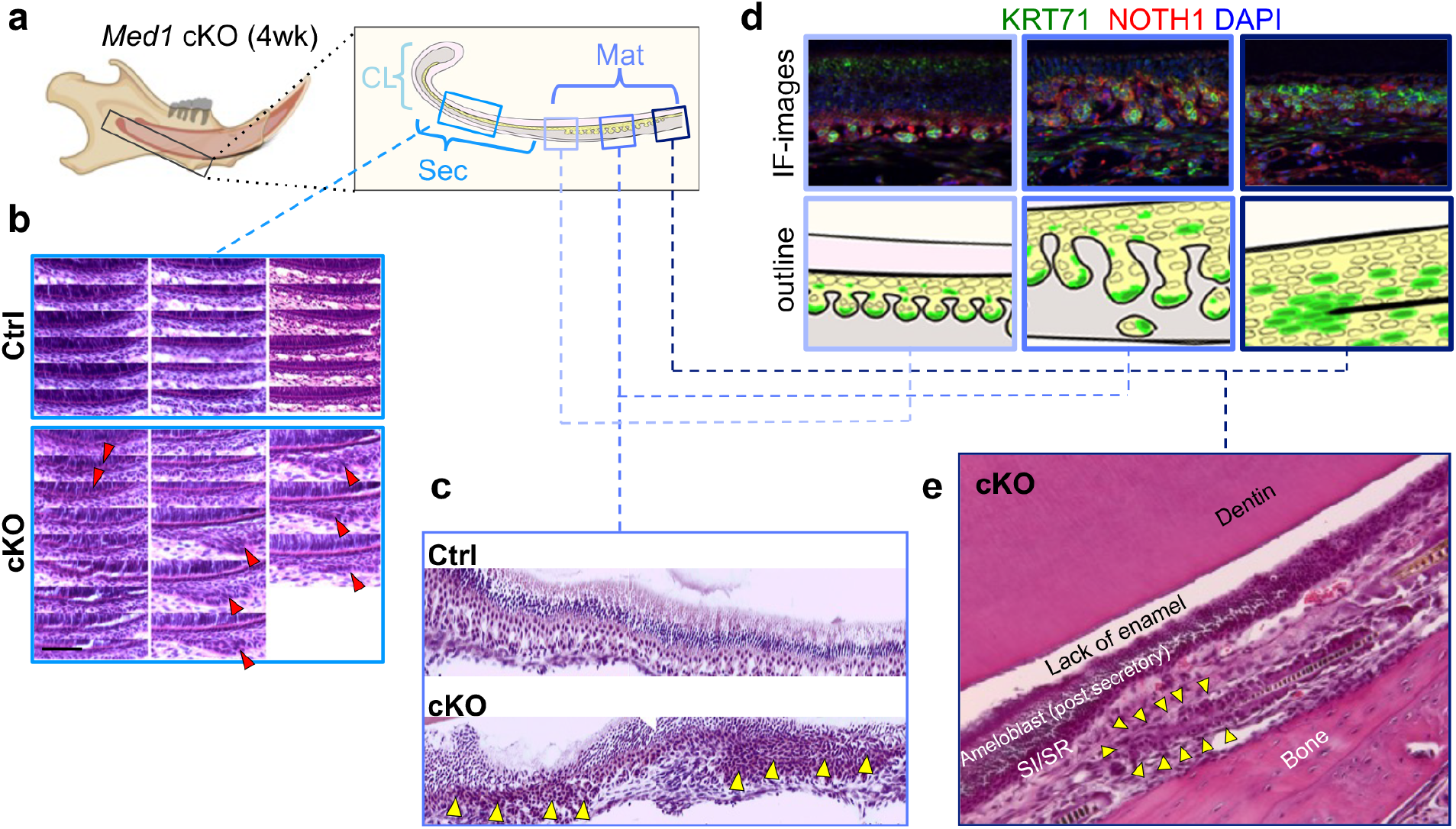
Histological and immuno-histological characterization of *Med1* cKO incisor tissue. **a** Diagram depicting the differentiation stages of mouse incisors including the location of the cervical loop (CL), the secretory stage (Sec) and the maturation stage (Mat) of dental epithelia. Right, detailed representation of Sec and Mat. **b** Serial sectioning showing progression of abnormal stratum intermedium (SI) development with generation of atypical cell clusters (red triangles) in *Med1* cKO mice. **c** H&E staining to show abnormal expansion of papillary layer in cKO (yellow triangles). **d** Top, immuno-fluorescent visualization of KRT17 (green), NOTCH1 (red) and DAPI counterstaining (blue) in SI/stellate reticulum (SR) derived papillary layers at 3 different locations of the Mat stage in *Med1* cKO mice. Bottom, diagram to show the locations of hair marker expression (green). **e** H&E-stained sections emphasizing the location of hair generating cell cluster found between dentin and bone in *Med1* cKO. SI, stratum intermedium, SR stellate reticulum.

**Suppl. Fig. 2:**
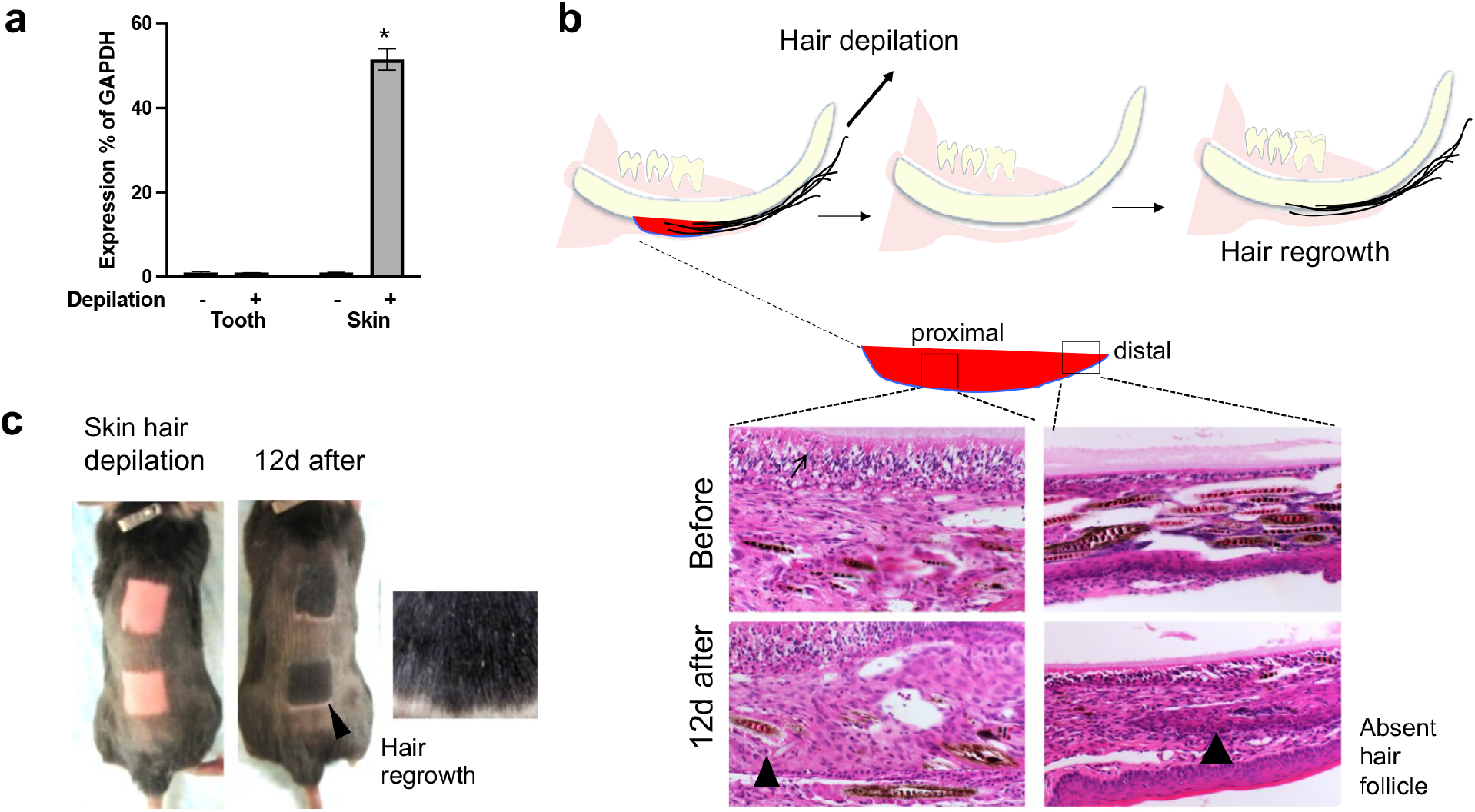
Effects of hair depilation on dental hair regeneration. **a** Hair marker *Krt31* mRNA expression in CL tissues of *Med1* cKO mice (tooth, left) and in normal skin tissue (skin right) before (-) and 8 days after (+) hair depilation. Representative data of average and SD of relative expression (% of GAPDH) and the statistical significance (n=3, * p<0.05). **b** Histological assessment of atypical hair generating cell clusters in *Med1* cKO incisors before and 12 days after hair depilation in proximal and distal region of the mandible (black triangles) corresponding to diagram. **c** Mouse skin shortly after hair depilation (left) and 12 days post depilation (right), when hair is regrown (arrow) and enlarged image of hair.

**Suppl. Fig. 3:**
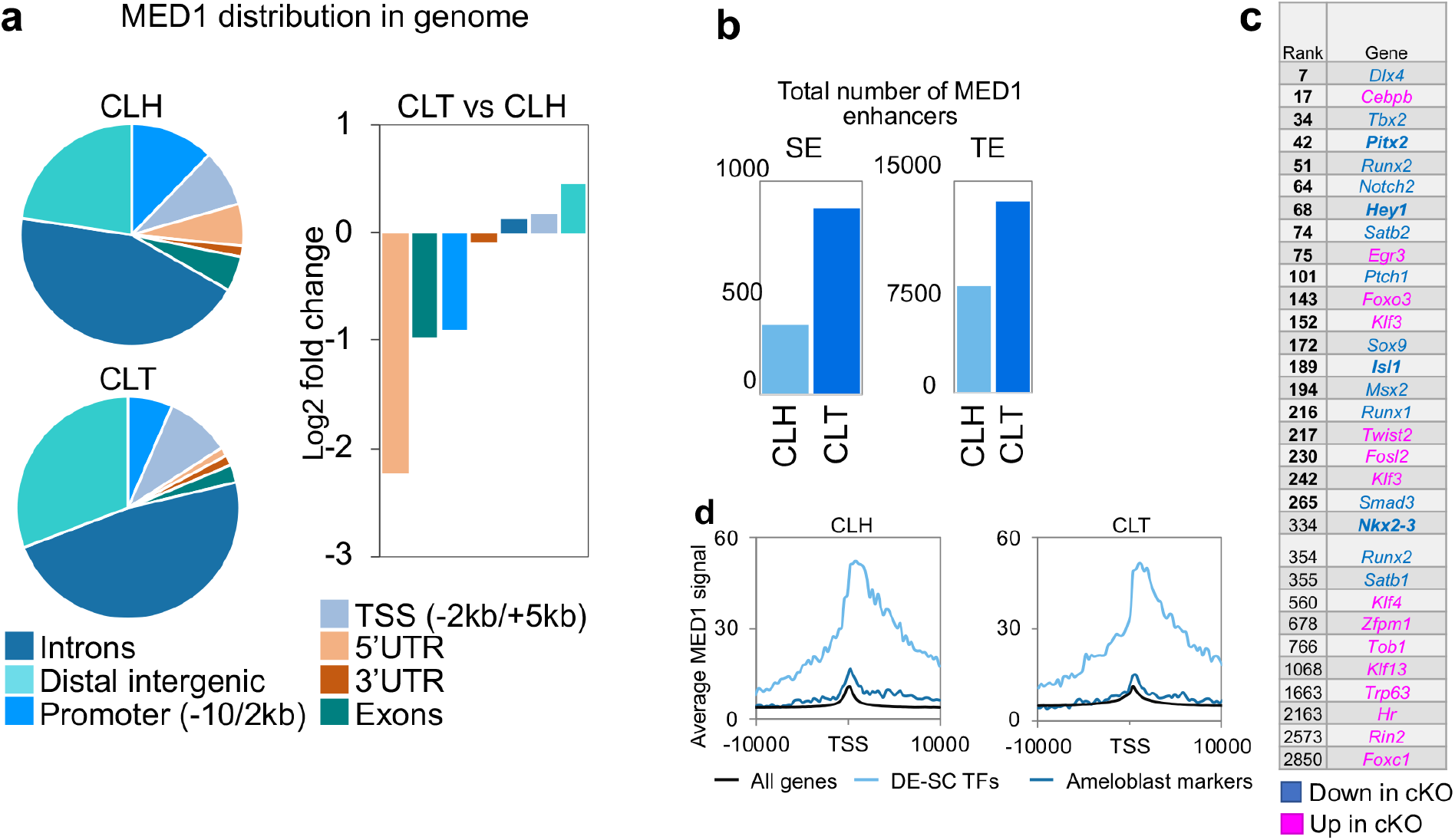
Med1 is distributed in different genomic regions in dental stem cells (CLH) and their progenies (CLT) in normal mice. **a** Left, relative distribution of Med1 binding at different genomic elements in CLH and CLT Ctrl tissues (left); right, shift in Med1 binding at shown elements between CLT and CLH of Ctrl tissues (right) **b** Total number of super enhancers (SE) and typical enhancers (TE) in CLH and CLT Ctrl tissues. The 325 SEs and 871 SEs were identified in the CLH and CLT, respectively. **c** List of transcription factors regulated by Med1 containing enhancers in CLH. Enhancer ranking numbers is noted left and colors define regulation of mRNA expression in CLH upon loss of *Med1*. **d** Average Med1 occupancy at promoters (TSS +/- 10kbp) for enamel fate transcription factors (TFs) (pale blue line), ameloblast markers (blue line) and all other genes (black line) in CLH and CLT tissues.

**Suppl. Fig. 4:**
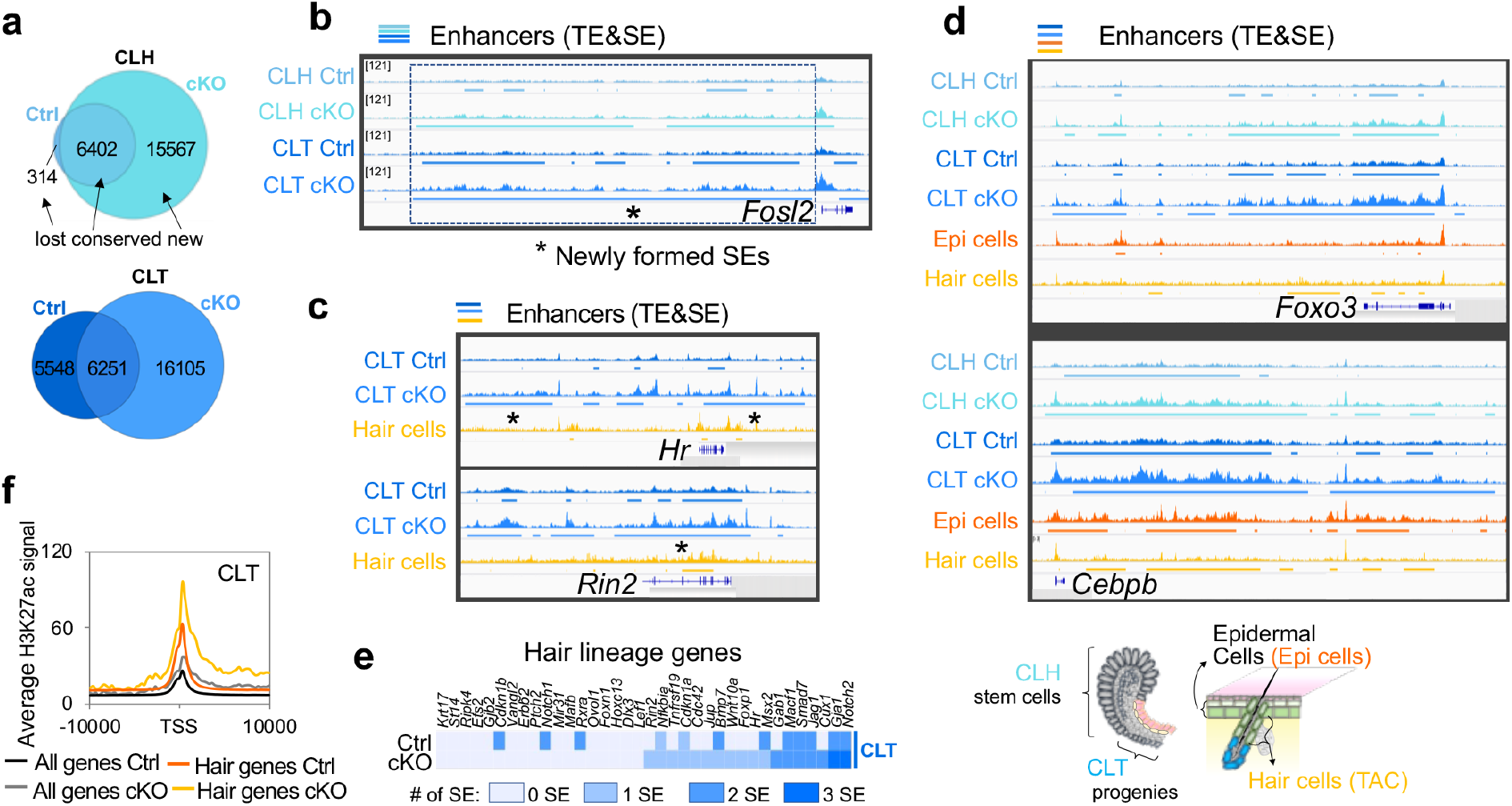
Genomic H3K27ac occupancy in promoters and enhancers and their changes in cKO dental tissues. **a** Numbers of typical enhancers (TEs) in Ctrl and cKO at CLH and CLT tissues. Lost, conserved and new TEs for cKO are shown. **b-d** H3K27ac occupancy around the loci of epidermal *Fosl2* (**b**), hair lineage driving *Hr* and *Rin2* (**c**), and epidermal transcription factors *Foxo3* and *Cebpb* (**d**) in Ctrl and *Med1* cKO at CLH and CLT tissues. Newly formed super-enhancers (SEs) in cKO are marked with asterisks (*). These enhancer profiles are compared with enhancer profiles (H3K27ac) from skin derived epidermal cells (Epi cells) and hair follicle transient amplifying (TAC) cells (Hair cells), in which locations of dental and skin epithelia are inserted. **e** Number of H3K27ac super-enhancers around genomic loci for hair lineage genes after *Med1* loss in CLT tissues. **f** Average H3K27ac occupancy at promoters (TSS +/- 10kbp) for hair lineage genes in CLT tissues in *Med1* cKO and Ctrl mice.

**Suppl. Fig. 5:**
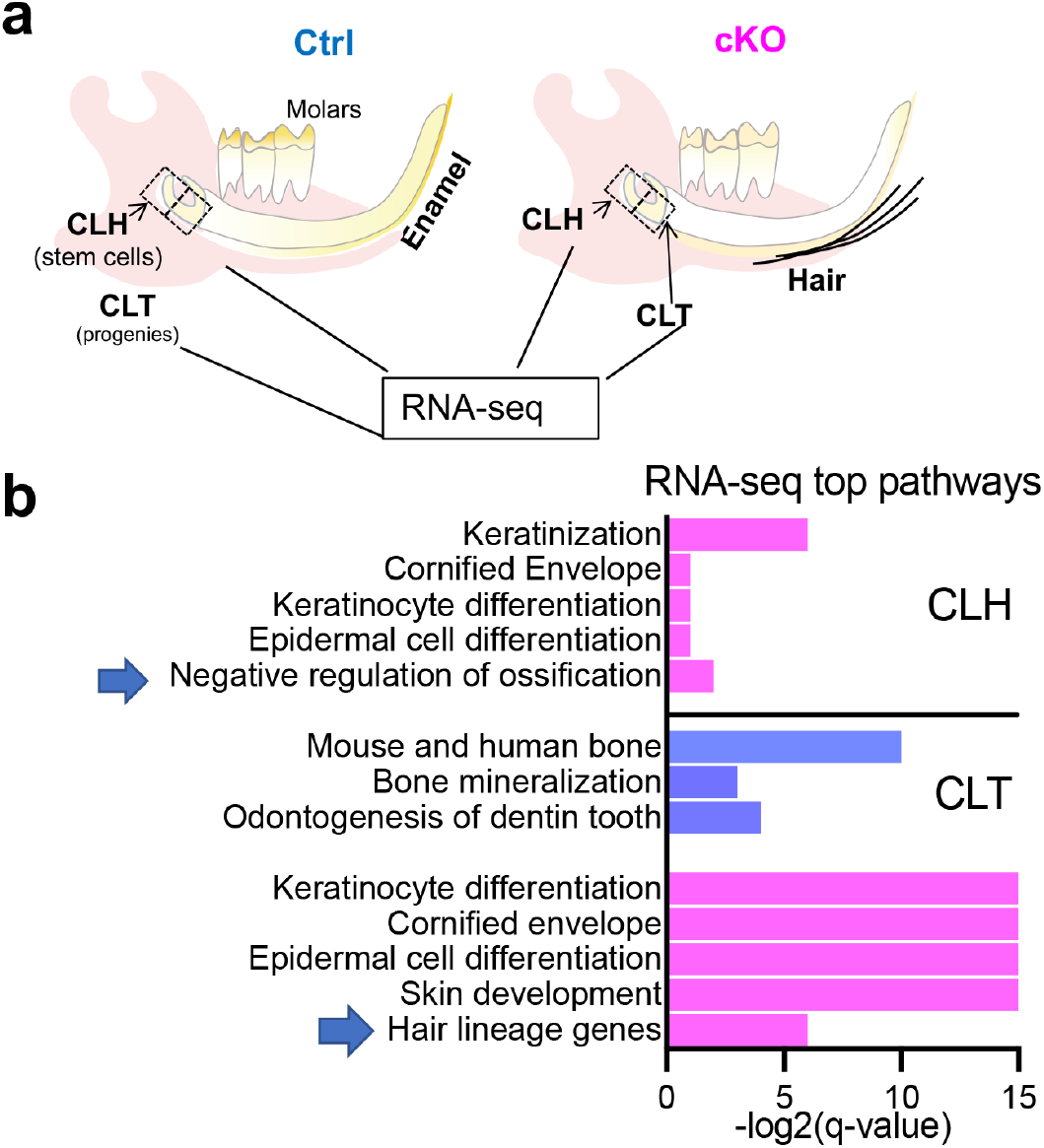
Functional annotation for differentially expressed genes in *Med1* cKO versus Ctrl dental stem cells (CLH) and their progenies (CLT). **a** Top, schematic diagram illustrating the collected tissues for the RNA-Seq experiment using two litters of 4 weeks-old Med1 cKO and Ctrl mice. **b** GO-based annotations representing up-regulated gene sets are shown in pink, down-regulated gene sets are shown in blue. Blue arrows show relevant pathways for hair formation (bottom) and enamel dysplasia (top) observed in *Med1* cKO mice.

**Suppl. Fig. 6:**
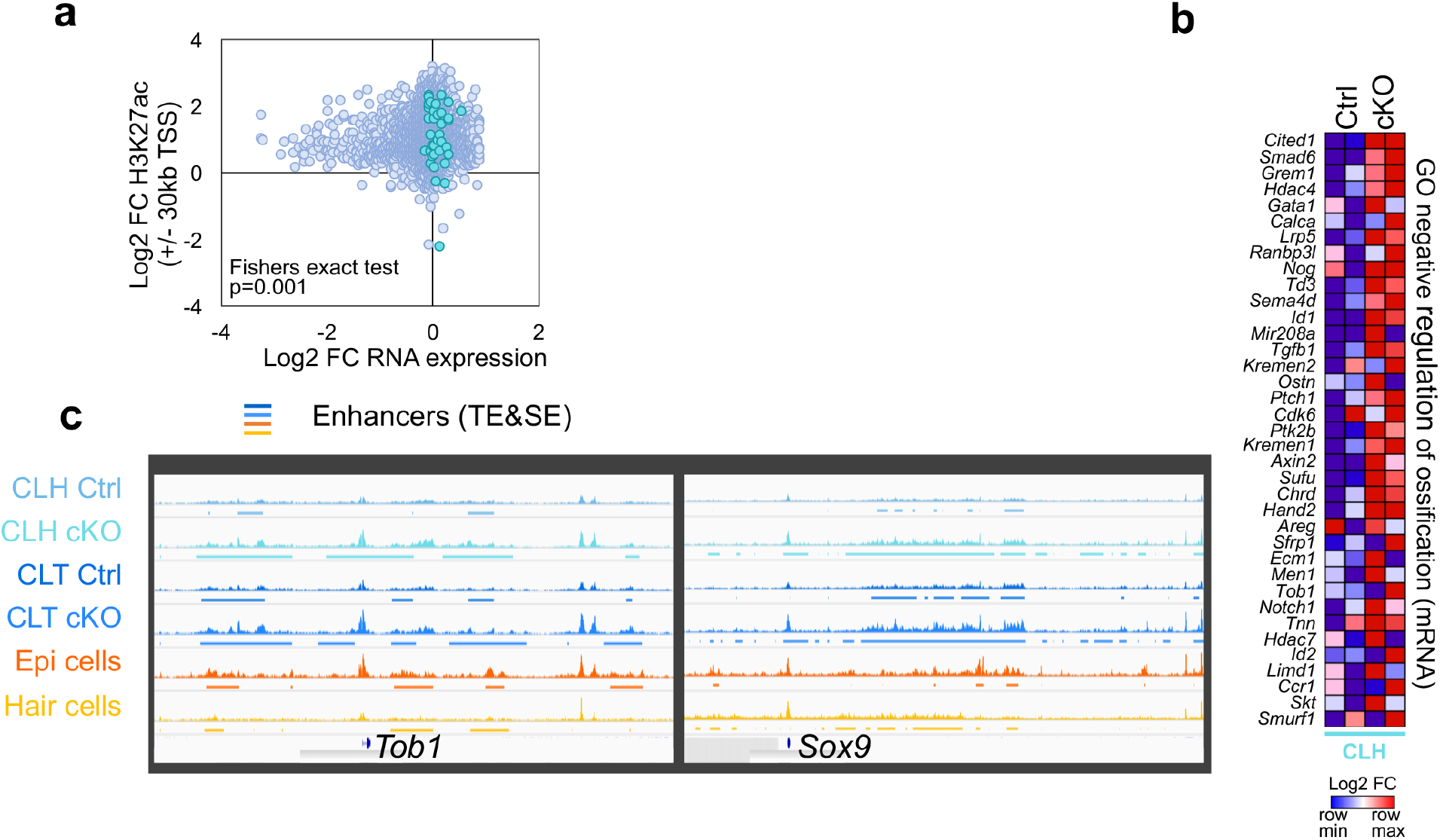
Loss of Med1 perturbs ossification pathways in dental tissues. **a** Correlation between gene expression and H3K27ac promoter occupancy (TSS +/-30 kb) in *Med1* cKO vs Ctrl CLT tissues. Blue dots highlight the gene-set ‘negative regulation of ossification’, pale blue dots represent all other genes. **b** Heatmap showing the changes is RNA expression for the gene set ‘Go negative regulation of ossification’ in Ctrl and *Med1* cKO CLH tissues. **c** Chip-seq enhancer profiles (H3K27ac) around loci of ossification inhibitors *Tob1 and Sox9*. Here, enhancers (colored underlines) expand in size or heights after loss of *Med1* cKO in CLH and CLT tissues. As a comparison, H3K27ac profiles for skin derived epidermal cells (Epi cells) and hair follicle TAC cells (Hair cells) are shown.

## REFERENCES

1 Wabik, A. & Jones, P. H. Switching roles: the functional plasticity of adult tissue stem cells. EMBO J 34, 1164–1179 (2015). https://doi.org:10.15252/embj.201490386

2 Garcia-Prat, L., Sousa-Victor, P. & Munoz-Canoves, P. Proteostatic and Metabolic Control of Stemness. Cell Stem Cell 20, 593–608 (2017). https://doi.org:10.1016/j.stem.2017.04.011

3 Middendorp, S. et al. Adult stem cells in the small intestine are intrinsically programmed with their location-specific function. Stem Cells 32, 1083–1091 (2014). https://doi.org:10.1002/stem.1655

4 Adam, R. C. et al. Pioneer factors govern super-enhancer dynamics in stem cell plasticity and lineage choice. Nature 521, 366–370 (2015). https://doi.org:10.1038/nature14289

5 Kaltschmidt, B., Kaltschmidt, C. & Widera, D. Adult craniofacial stem cells: sources and relation to the neural crest. Stem Cell Rev Rep 8, 658–671 (2012). https://doi.org:10.1007/s12015-011-9340-9

6 Jimenez-Rojo, L., Granchi, Z., Graf, D. & Mitsiadis, T. A. Stem Cell Fate Determination during Development and Regeneration of Ectodermal Organs. Front Physiol 3, 107 (2012). https://doi.org:10.3389/fphys.2012.00107

7 Yoshizaki, K., Fukumoto, S., Bikle, D. D. & Oda, Y. Transcriptional Regulation of Dental Epithelial Cell Fate. Int J Mol Sci 21 (2020). https://doi.org:10.3390/ijms21238952

8 Manini, I. et al. Multi-potent progenitors in freshly isolated and cultured human mesenchymal stem cells: a comparison between adipose and dermal tissue. Cell Tissue Res 344, 85–95 (2011). https://doi.org:10.1007/s00441-011-1139-0

9 Bianco, P. “Mesenchymal” stem cells. Annu Rev Cell Dev Biol 30, 677–704 (2014). https://doi.org:10.1146/annurev-cellbio-100913-013132

10 Bianco, P., Robey, P. G. & Simmons, P. J. Mesenchymal stem cells: revisiting history, concepts, and assays. Cell Stem Cell 2, 313–319 (2008). https://doi.org:10.1016/j.stem.2008.03.002

11 Wang, X. P. et al. An integrated gene regulatory network controls stem cell proliferation in teeth. PLoS Biol 5, e159 (2007). https://doi.org:10.1371/journal.pbio.0050159

12 Klein, O. D. et al. An FGF signaling loop sustains the generation of differentiated progeny from stem cells in mouse incisors. Development 135, 377–385 (2008). https://doi.org:10.1242/dev.015081

13 Mitsiadis, T. A. & Graf, D. Cell fate determination during tooth development and regeneration. Birth Defects Res C Embryo Today 87, 199–211 (2009). https://doi.org:10.1002/bdrc.20160

14 Fuchs, E., Tumbar, T. & Guasch, G. Socializing with the neighbors: stem cells and their niche. Cell 116, 769–778 (2004). https://doi.org:S0092867404002557 [pii]

15 Juuri, E. et al. Sox2 marks epithelial competence to generate teeth in mammals and reptiles. Development 140, 1424–1432 (2013). https://doi.org:10.1242/dev.089599

16 Juuri, E. et al. Sox2+ stem cells contribute to all epithelial lineages of the tooth via Sfrp5+ progenitors. Dev Cell 23, 317–328 (2012). https://doi.org:10.1016/j.devcel.2012.05.012

17 Morita, R. et al. Tracing the origin of hair follicle stem cells. Nature 594, 547–552 (2021). https://doi.org:10.1038/s41586-021-03638-5

18 Gonzales, K. A. U. & Fuchs, E. Skin and Its Regenerative Powers: An Alliance between Stem Cells and Their Niche. Dev Cell 43, 387–401 (2017). https://doi.org:10.1016/j.devcel.2017.10.001

19 Sundberg, J. P., Peters, E. M. & Paus, R. Analysis of hair follicles in mutant laboratory mice. J Investig Dermatol Symp Proc 10, 264–270 (2005). https://doi.org:10.1111/j.1087-0024.2005.10126.x

20 Botchkarev, V. A. & Flores, E. R. p53/p63/p73 in the epidermis in health and disease. Cold Spring Harb Perspect Med 4 (2014). https://doi.org:10.1101/cshperspect.a015248

21 Greco, V. et al. A two-step mechanism for stem cell activation during hair regeneration. Cell Stem Cell 4, 155–169 (2009). https://doi.org:10.1016/j.stem.2008.12.009

22 Alonso, L. & Fuchs, E. The hair cycle. J Cell Sci 119, 391–393 (2006). https://doi.org:10.1242/jcs.02793

23 Sakai, Y. & Demay, M. B. Evaluation of keratinocyte proliferation and differentiation in vitamin D receptor knockout mice. Endocrinology 141, 2043–2049 (2000).

24 Lin, C. R. et al. Pitx2 regulates lung asymmetry, cardiac positioning and pituitary and tooth morphogenesis. Nature 401, 279–282 (1999). https://doi.org:10.1038/45803

25 Han, X. et al. The transcription factor NKX2-3 mediates p21 expression and ectodysplasin-A signaling in the enamel knot for cusp formation in tooth development. J Biol Chem 293, 14572–14584 (2018). https://doi.org:10.1074/jbc.RA118.003373

26 Naveau, A. et al. Isl1 Controls Patterning and Mineralization of Enamel in the Continuously Renewing Mouse Incisor. J Bone Miner Res 32, 2219–2231 (2017). https://doi.org:10.1002/jbmr.3202

27 Panteleyev, A. A., Paus, R., Ahmad, W., Sundberg, J. P. & Christiano, A. M. Molecular and functional aspects of the hairless (hr) gene in laboratory rodents and humans. Exp Dermatol 7, 249–267 (1998). https://doi.org:10.1111/j.1600-0625.1998.tb00295.x-i1

28 Basel-Vanagaite, L. et al. RIN2 deficiency results in macrocephaly, alopecia, cutis laxa, and scoliosis: MACS syndrome. Am J Hum Genet 85, 254–263 (2009). https://doi.org:10.1016/j.ajhg.2009.07.001

29 Whyte, W. A. et al. Master transcription factors and mediator establish super-enhancers at key cell identity genes. Cell 153, 307–319 (2013). https://doi.org:10.1016/j.cell.2013.03.035

30 Malik, S. & Roeder, R. G. Dynamic regulation of pol II transcription by the mammalian Mediator complex. Trends Biochem Sci 30, 256–263 (2005). https://doi.org:S0968-0004(05)00081-2 [pii] 10.1016/j.tibs.2005.03.009

31 Kornberg, R. D. Mediator and the mechanism of transcriptional activation. Trends Biochem Sci 30, 235–239 (2005).

32 Taatjes, D. J. The human Mediator complex: a versatile, genome-wide regulator of transcription. Trends Biochem Sci 35, 315–322 (2010). https://doi.org:S0968-0004(10)00025-3 [pii] 10.1016/j.tibs.2010.02.004

33 Oda, Y. et al. Coactivator MED1 ablation in keratinocytes results in hair-cycling defects and epidermal alterations. J Invest Dermatol 132, 1075–1083 (2012). https://doi.org:10.1038/jid.2011.430

34 Oda, Y., Chalkley, R. J., Burlingame, A. L. & Bikle, D. D. The transcriptional coactivator DRIP/mediator complex is involved in vitamin D receptor function and regulates keratinocyte proliferation and differentiation. J Invest Dermatol 130, 2377–2388 (2010). https://doi.org:10.1038/jid.2010.148

35 Yue, X., Izcue, A. & Borggrefe, T. Essential role of Mediator subunit Med1 in invariant natural killer T-cell development. Proc Natl Acad Sci U S A 108, 17105–17110 (2011). https://doi.org:10.1073/pnas.1109095108

36 Jiang, P. et al. Key roles for MED1 LxxLL motifs in pubertal mammary gland development and luminal-cell differentiation. Proc Natl Acad Sci U S A 107, 6765–6770 (2010). https://doi.org:10.1073/pnas.1001814107

37 Jia, Y. et al. Peroxisome proliferator-activated receptor-binding protein null mutation results in defective mammary gland development. J Biol Chem 280, 10766–10773 (2005). https://doi.org:10.1074/jbc.M413331200

38 Zhang, X. et al. MED1/TRAP220 exists predominantly in a TRAP/ Mediator subpopulation enriched in RNA polymerase II and is required for ER-mediated transcription. Mol Cell 19, 89–100 (2005). https://doi.org:10.1016/j.molcel.2005.05.015

39 Yoshizaki, K. et al. Ablation of coactivator Med1 switches the cell fate of dental epithelia to that generating hair. PLoS One 9, e99991 (2014). https://doi.org:10.1371/journal.pone.0099991

40 Yoshizaki, K. et al. Mediator 1 contributes to enamel mineralization as a coactivator for Notch1 signaling and stimulates transcription of the alkaline phosphatase gene. J Biol Chem 292, 13531–13540 (2017). https://doi.org:10.1074/jbc.M117.780866

41 Xuan, D. et al. Effect of cleidocranial dysplasia-related novel mutation of RUNX2 on characteristics of dental pulp cells and tooth development. J Cell Biochem 111, 1473–1481 (2010). https://doi.org:10.1002/jcb.22875

42 Chu, Q. et al. Ablation of Runx2 in Ameloblasts Suppresses Enamel Maturation in Tooth Development. Sci Rep 8, 9594 (2018). https://doi.org:10.1038/s41598-018-27873-5

43 Zhang, Y. et al. SATB1 establishes ameloblast cell polarity and regulates directional amelogenin secretion for enamel formation. BMC Biol 17, 104 (2019). https://doi.org:10.1186/s12915-019-0722-9

44 Hnisz, D. et al. Super-enhancers in the control of cell identity and disease. Cell 155, 934–947 (2013). https://doi.org:10.1016/j.cell.2013.09.053

45 Lien, W. H. et al. Genome-wide maps of histone modifications unwind in vivo chromatin states of the hair follicle lineage. Cell Stem Cell 9, 219–232 (2011). https://doi.org:10.1016/j.stem.2011.07.015

46 Wei, Y. et al. Human skeletal muscle-derived stem cells retain stem cell properties after expansion in myosphere culture. Exp Cell Res 317, 1016–1027 (2011). https://doi.org:10.1016/j.yexcr.2011.01.019

47 Kim, K. H. et al. EGR3 Is a Late Epidermal Differentiation Regulator that Establishes the Skin-Specific Gene Network. J Invest Dermatol 139, 615–625 (2019). https://doi.org:10.1016/j.jid.2018.09.019

48 House, J. S., Zhu, S., Ranjan, R., Linder, K. & Smart, R. C. C/EBPalpha and C/EBPbeta are required for Sebocyte differentiation and stratified squamous differentiation in adult mouse skin. PLoS One 5, e9837 (2010). https://doi.org:10.1371/journal.pone.0009837

49 Zhang, W. & Yelick, P. C. Tooth Repair and Regeneration: Potential of Dental Stem Cells. Trends Mol Med 27, 501–511 (2021). https://doi.org:10.1016/j.molmed.2021.02.005

50 Zheng, L. W., Linthicum, L., DenBesten, P. K. & Zhang, Y. The similarity between human embryonic stem cell-derived epithelial cells and ameloblast-lineage cells. Int J Oral Sci 5, 1–6 (2013). https://doi.org:10.1038/ijos.2013.14

51 Takeo, M. et al. Expansion and characterization of epithelial stem cells with potential for cyclical hair regeneration. Sci Rep 11, 1173 (2021). https://doi.org:10.1038/s41598-020-80624-3

52 Hu, X. et al. Efficient induction of functional ameloblasts from human keratinocyte stem cells. Stem Cell Res Ther 9, 126 (2018). https://doi.org:10.1186/s13287-018-0822-4

53 Jia, Y. et al. Transcription coactivator PBP, the peroxisome proliferator-activated receptor (PPAR)-binding protein, is required for PPARalpha-regulated gene expression in liver. J Biol Chem 279, 24427–24434 (2004). https://doi.org:10.1074/jbc.M402391200 M402391200 [pii]

